# A novel telencephalon-opto-hypothalamic morphogenetic domain produces most of the glutamatergic neurons of the medial extended amygdala

**DOI:** 10.1101/2020.07.17.207936

**Authors:** Lorena Morales, Beatriz Castro-Robles, Antonio Abellán, Ester Desfilis, Loreta Medina

## Abstract

Deficits in social cognition and behavior are a hallmark of many psychiatric disorders. The medial extended amygdala, including the medial amygdala and the medial bed nucleus of the stria terminalis, is a key component of functional networks involved in sociality. However, this nuclear complex is highly heterogeneous and contains numerous GABAergic and glutamatergic neuron subpopulations. Deciphering the connections of different neurons is essential in order to understand how this structure regulates different aspects of sociality, and it is necessary to evaluate their differential implication in distinct mental disorders. Developmental studies in different vertebrates are offering new venues to understand neuronal diversity of the medial extended amygdala, and are helping to establish a relation between the embryonic origin and molecular signature of distinct neurons with the functional subcircuits in which they are engaged. These studies have provided many details on the distinct GABAergic neurons of the medial extended amygdala, but information on the glutamatergic neurons is still scarce. Using an Otp-eGFP transgenic mouse and multiple fluorescent labeling, we show that most glutamatergic neurons of the medial extended amygdala originate in a novel telencephalo-opto-hypothalamic embryonic domain (TOH), located at the transition between telencephalon and hypothalamus, which produces Otp-lineage neurons expressing the telencephalic marker Foxg1 but not Nkx2.1 during development. These glutamatergic cells include a subpopulation of projection neurons of the medial amygdala, which activation has been previously shown to promote autistic-like behavior. Our data open new venues for studying the implication of this neuron subtype in neurodevelopmental disorders producing social deficits.

## Introduction

The medial amygdala and medial part of the bed nucleus of the stria terminalis (BSTM) form part of a cellular continuum of the ventral telencephalon, known as the ‘medial extended amygdala’ (Alheid and Heimer, 1988; de Olmos et al., 2004), which plays a critical role in social cognition and control of socio-sexual behaviors (Swanson and Petrovich, 1989; Martínez-García et al. 2002; 2007; Choi et al., 2005; Phelps and LeDoux, 2005; Bickart et al., 2014; Medina et al., 2011, 2017a, 2019; Pessoa et al., 2019). Although located in the ventral telencephalon, these nuclei contain not only abundant GABAergic neurons (as typical of the subpallium; Swanson and Petrovich, 1989; McDonald, 2003), but also an important population of glutamatergic neurons (Choi et al., 2005). Both GABAergic and glutamatergic neurons of the medial extended amygdala are heterogeneous, including different subtypes, and each subtype appears to be involved in a specific functional pathway (Choi et al., 2005; Kiss et al., 2011). By way of cell-specific functional manipulation using optogenetics, GABAergic and glutamatergic neurons of the medial amygdala have been shown to exert antagonistic influences on social behavior: while GABAergic neurons promote different types of social interactions, glutamatergic neurons suppress social behavior and promote self-grooming, resembling autistic-like features (Hong et al., 2014).

Developmental studies have provided an explanation for the presence of different subtypes of GABAergic and glutamatergic neurons in the medial extended amygdala: these different neurons originate in distinct embryonic domains of the pallium, subpallium, hypothalamus and prethalamic eminence (García-López et al., 2008; Soma et al., 2009; Hirata et al., 2009; Carney et al., 2010; García-Moreno et al., 2010; Bupesh et al., 2011a,b; Lischinsky et al., 2017; Medina et al., 2011, 2017a; Ruiz-Reig et al., 2018). Neurons with different embryonic origin express distinct combinatorial expression of transcription factors during development, and appear to be engaged in different functional pathways (García-López et al., 2008; Hirata et al., 2009; Carney et al., 2010; Medina et al., 2011; Sokolowski and Corbin, 2012; Abellán et al., 2013; Lischinsky et al., 2017). This has been studied in more detail for the GABAergic neurons of the medial extended amygdala, which derive from either progenitors of the ventrocaudal pallidum expressing Nkx2.1 and Lhx6, or progenitors of the preoptic area expressing Nkx2.1 and Shh (García-López et al., 2008; Carney et al., 2010). GABAergic neurons of the medial extended amygdala with origin in the ventrocaudal pallidal division (also called diagonal domain by Puelles et al., 2016) retain expression of Lhx6 during ontogenesis and become connected with Lhx6-rich areas of the preoptic region and ventromedial hypothalamus related to sexual behavior (García-López et al., 2008; Medina et al., 2011; Sokolowski and Corbin, 2012; Abellán et al., 2013). On the contrary, GABAergic neurons of the medial amygdala derived from the preoptic area express Shh (García-López et al., 2008; Carney et al., 2010; Medina et al., 2011; Abellán et al., 2013), Dbx1 (Hirata et al., 2009) or Foxp2 (Lischinsky et al., 2017), and are involved in different types of social interactions, such as mating and aggression (Lischinsky et al., 2017). Regarding the glutamatergic neurons, these appear to derive from the ventral/ventrocaudal pallium (expressing Lhx9 and/or Ebf3; ventral pallium: García-López et al., 2008; Bupesh et al., 2011a; Medina et al., 2011; ventrocaudal pallium: Medina et al., 2017a; Ruiz-Reig et al., 2018) or from domains outside the telencephalon, including the supraopto-paraventricular domain of the hypothalamus (Otp-expressing) and the prethalamic eminence (Lhx5-expressing) (Soma et al., 2009; García-Moreno et al., 2010; Bupesh et al., 2011a; discussed by Medina et al., 2011, 2017; Abellán et al., 2013). However, the exact contribution and the specific phenotype of neurons from each different domain reaching the medial extended amygdala are unknown. This information could contribute to understand the different subcircuits and functions in which different glutamatergic cells are engaged.

Foxg1 is one of the earliest transcription factors expressed in the telencephalic anlage during development, and its inactivation produces proliferation defects in telencephalic progenitors, which leads to severe telencephalic hypoplasia, with complete absence of the subpallium (Xuan et al., 1995; Dou et al., 1999; Martynoga et al., 2005; Manuel et al., 2010, 2011). After this initial period, telencephalic postmitotic cells continue to express Foxg1 throughout embryonic, fetal and postnatal development (Dou et al. 1999; Manuel et al., 2011). Based on this, Foxg1 is often considered a good marker to distinguish between telencephalic and non-telencephalic regions (for example, Carney et al., 2010). However, a recent study in zebrafish has shown the existence of a new domain at the transition between telencephalon and hypothalamus that expresses both Otp and Foxg1, which was named the ‘optic recess region’ because of its relation to part of the eye vesicle (Affaticati et al., 2015). This raises questions on the existence of such domain in other vertebrates, including mammals, and - if present - on the contribution of this domain to the formation of telencephalic structures such as the medial extended amygdala. Thus, the aim of this study was to investigate these issues in mouse. To overcome the problem that Otp downregulates its expression in many neurons during middle stages of development, we took advantage of Otp-eGFP transgenic mice, with permanent labeling of Otp-lineage cells throughout ontogenesis. Embryonic and postnatal brains were processed for multiple fluorescence labeling to analyze co-expression of GFP (Otp-lineage) with Foxg1, other region-specific transcription factors, radial glial fibers, VGLUT2 (for glutamatergic cells) and GAD67 (for GABAergic cells). Our results showed the existence in mouse of a new radial histogenetic domain at the frontier between telencephalon and hypothalamus, which produces cells expressing both Otp and Foxg1. This new radial unit produces the majority of the glutamatergic neurons of the medial extended amygdala, the preoptic area and the dorsal (anterior) paraventricular hypothalamic region. Although this unit resembles that previously described in zebrafish (Affaticati et al., 2015), this division is not only related to part of the optic recess, but most importantly to parts of the telencephalon and adjacent hypothalamic region, implying a redefinition of the telencephalon-hypothalamic boundary. For this reason, we have called this division the telencephalo-opto-hypothalamic transition domain (TOH).

## Material and Methods

In the present study, we used wild type mice (*Mus musculus*, Swiss) and Otp-eGFP transgenic mice (*Mus musculus*, Tg (Otp-EGFP) OI121Gsat/Mmucd; Mutant Mouse Resource & Research Centers, MMRRC supported by NIH, University of California at Davis, USA), from embryonic day 12.5 (post-coitum) (E12.5) to postnatal day 19 (P19). The sex of animals was determined by a method based on PCR (McClive and Sinclair, 2001). Progenitors and weaned-off postnatal animals were housed in groups of three to five at 22 ± 2°C on a 12-h light/dark cycle, with food and water ad libitum, in the rodent animal facility of the University of Lleida (REGA license no. ES251200037660). Wild type animals were in the conventional area, while transgenic mice were kept in the pathogen-free area, which fulfills all requirements for genetically-modified animals (notification no. A/ES/19/I-06). All the animals were treated according to the regulations and laws of the European Union (Directive 2010/63/EU) and the Spanish Government (Royal Decrees 53/2013) for the care and handling of animals in research. All the protocols used were approved by the Committees of Ethics for Animal Experimentation and Biosecurity of the University of Lleida.

### Tissue collection and fixation

At appropriate development days, the mouse embryos were obtained by cesarean section from pregnant females, which were previously sacrificed by cervical dislocation or by a lethal dose of sodium pentobarbital (0.1mg/g; i.p.). Upon extraction, early embryos (E12.5-E14.5) were rapidly sacrificed by decapitation. The brains were dissected and fixed by immersion in phosphate-buffered 4% paraformaldehyde (4% PFA; pH7.4; 0.1M) overnight at 4°C. Older embryos and postnatal animals (E16.5 to P19) were deeply anesthetized with sodium pentobarbital (0.1mg/g; i.p.) and then transcardially perfused with 0.9% saline solution (0.9% NaCl), followed by 4% PFA. After dissection, the brains were postfixed by immersion in 4% PFA overnight at 4°C.

### Sample preparation and sectioning

Some brains were embedded in 4% low-melt agarose (LOW EEO, Laboratorios Conda S.A., Spain) in 0,1 M phosphate-buffered saline (PBS) and sectioned using a vibratome (Leica VT 1000S, Leica Microsystems GmbH, Germany) on the frontal, sagittal, horizontal or oblique-horizontal plane at 80 µm-thick. Other brains were cryoprotected with 30% sucrose in phosphate buffer (PB; 0.1M; pH 7.4) or with a solution of glycerol and DMSO in PB and were frozen with −60/-70°C isopentane (2-methyl butane, Sigma-Aldrich, Germany) (following the protocol of Rosene et al, 1986). The frozen brains were preserved at −80°C until use. Frontal, horizontal or sagittal free-floating sections were obtained using a freezing sliding microtome (60 µm-thick, Microm HM 450, Thermo Fisher Scientific, United Kingdom) or a cryostat (40 µm-thick, Leica CM 3000, Leica Microsystems GmbH). All sections were collected in 4°C PBS and then processed for in situ hybridization, immunohistochemistry or immunofluorescence.

**Figure 1.**
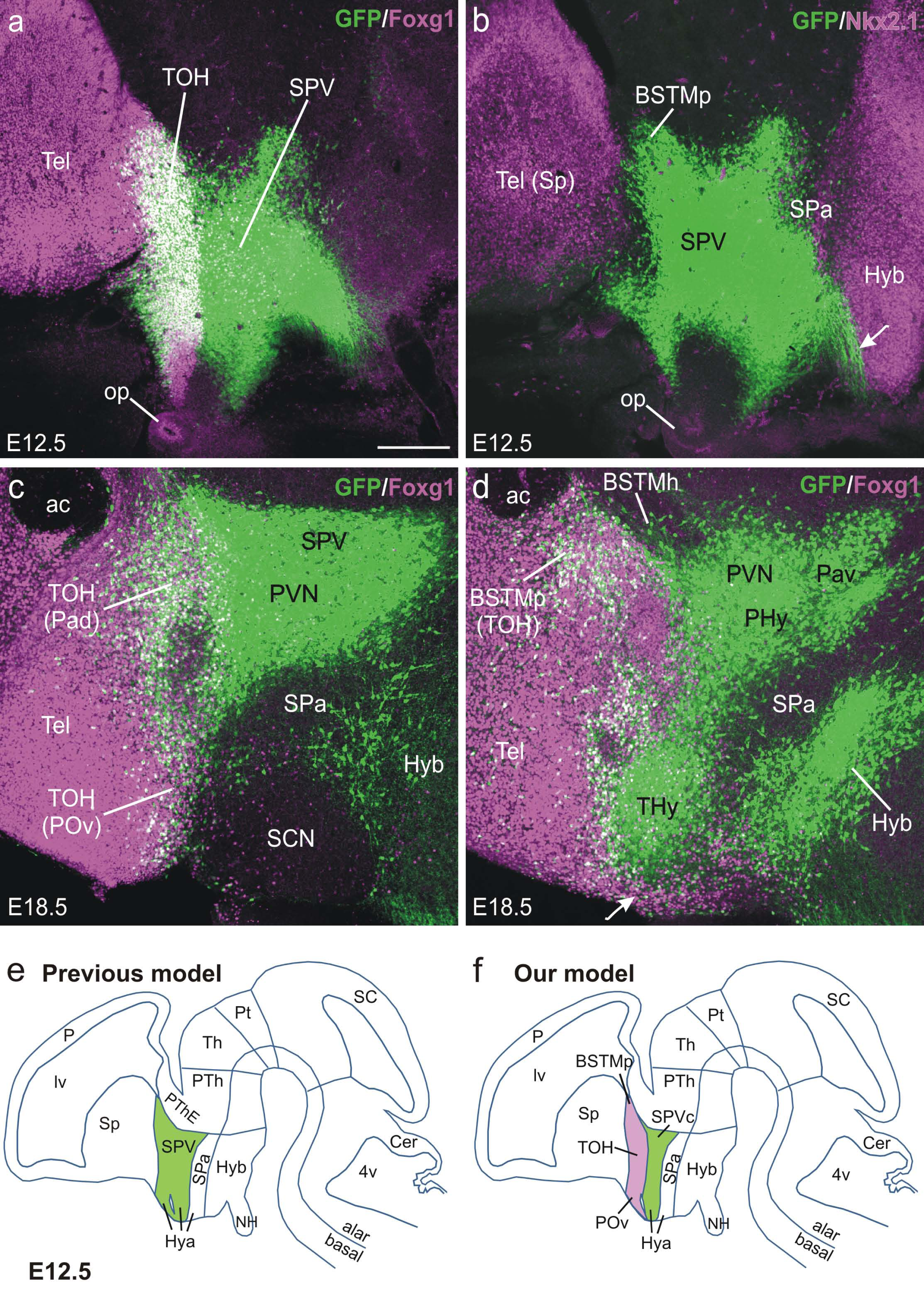
**a-d:** Images of sagittal brain sections of Otp-eGFP mice, at E12.5 (a,b) and E18.5 (c,d), double labeled for GFP (green) and Foxg1 (magenta). GFP is expressed in the classical supraopto-paraventricular domain (SPV), but a part of it co-expresses the telencephalic marker Foxg1. This overlapping subdivision is here defined as the telencephalon-opto-hypothalamic domain (TOH). Schemes of E12.5 sagittal brain sections are shown in **e** and **f**, representing classical SPV versus its two subdivisions, here called TOH and SPV core. Arrow in d points to a group of Foxg1-expressing cells that forms a continuum between telencephalon and hypothalamus. See text for more details. For abbreviations see list. Scale bar in a = 200 µm (applies to a-d).

### Immunohistochemistry

Free floating brain sections of wild type embryos were processed for immunohistochemistry to detect Islet1/2, Otp or RC2 (a radial glial marker), while brain sections of Otp-eGFP embryos and postnatal animals were processed for immunohistochemistry to detect GFP.

Islet1/2, Otp and RC2: Some series of sections (those for Islet1) were initially treated with a citrate buffered solution with 0.05% Tween-20 (pH = 6) at 80°C for 20 minutes, a procedure for antigen retrieval. After washing, the rest of the procedure followed the regular protocol briefly described next. Sections were processed to inhibit the endogenous peroxidase activity by an incubation in 1% H_2_O_2_ in PBS during 10 minutes. Then, the tissue was permeabilized by washing with PBS containing 0.3% Triton X 100 (PBS-Tx; pH = 7,4; 0.1M), followed by an incubation with a blocking solution, containing 10% normal goat serum (NGS) and 2% of bovine serum albumin (BSA) in PBS-Tx, for 1 hour at room temperature. Then, the sections were incubated in the primary antibody (Table 1), diluted in PBS-Tx with 10% of normal goat serum (NGS; Vector Laboratories Ltd., United Kingdom), for 72 h at 4°C and gentle agitation.

**Table 1.**
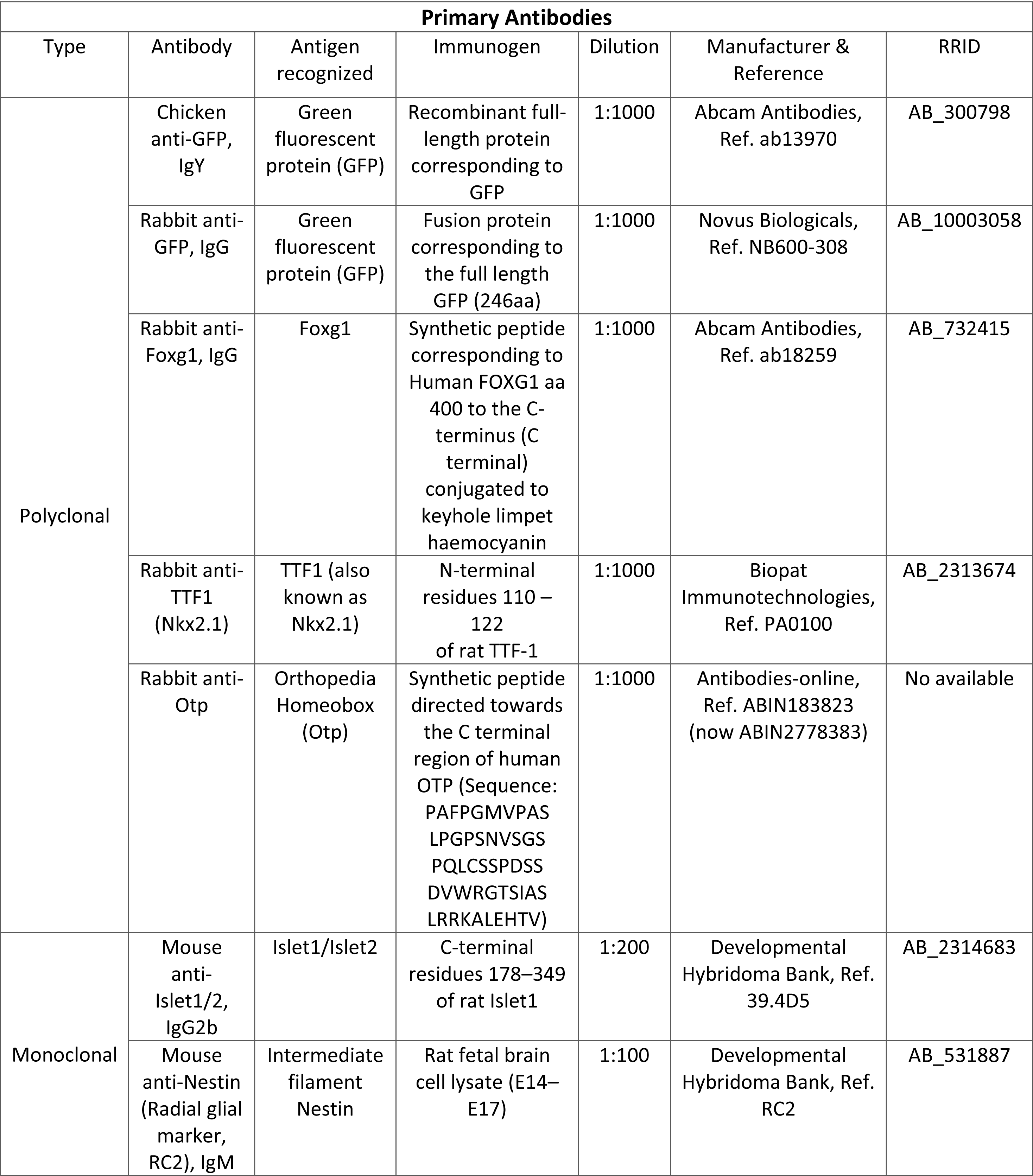
Primary antibodies.

After incubation in the primary antibody, the sections were rinsed in PBS and then incubated in a biotinylated secondary antibody (Table 2) diluted in PBS-Tx, for 90 minutes at room temperature and under gentle agitation. Following, the sections were washed and incubated with the avidin-biotin complex (AB Complex, Vector Laboratories Ltd.) for 1 hour at room temperature. After, the section were rinsed two times in PBS and then with a Tris buffer (0.05M, pH 7.6). Following this, the sections were incubated with diaminobenzidine (DAB), diluted in a Tris-buffered solution also containing urea and H_2_O_2_. Finally, the sections were rinsed and mounted with glycerol gelatin (Sigma-Aldrich, Germany). GFP: The free-floating sections were permeabilized by washing 3 times for 10 minutes with PBS-Tx, followed by an incubation with the blocking solution, for 1 hour at room temperature. Subsequently, the sections were incubated in a primary chicken anti-GFP antibody (Table 1) diluted in PBS-Tx for 72 hours at 4°C. After the washings in PBS-Tx, the endogenous peroxidase activity was reduced by an incubation in PBS 1% H_2_O_2_ and 10% methanol during 30 minutes. Then, the sections were washed and incubated in a biotinylated goat anti-chicken secondary antibody (Table 2), diluted in PBS-Tx, for 90-120 minutes at room temperature. Finally, the sections were washed and stained using the DAB method (as described above). Finally, the sections were washed, mounted using 0.25% gelatin in Tris buffer (TB; pH 8; 0.1M), dehydrated and coverslipped with Permount (Thermo Fisher Scientific).

**Table 2.** Secondary antibodies.

| **Secondary Antibodies** |  |  |  |
| --- | --- | --- | --- |
| Type | Antibody | Dilution | Manufacturer & Reference |
| **Biotinylated** | Goat anti-chicken IgY (H+L), biotinylated | 1:200 | Vector, Ref. BA-9010 |
|  | Goat anti-rabbit IgG, biotinylated | 1:200 | Vector, Ref BA-1000 |
|  | Goat anti-mouse IgG, biotinylated | 1:200 | Vector, BA-9200 |
|  | Goat anti-mouse IgM, biotinylated | 1:200 | Vector, BA-2020 |
| **Fluorescent** | Goat anti-chicken IgY (H+L), coupled to Alexa 488 | 1:500 | Invitrogen, Ref. A-11039 |
|  | Donkey anti-rabbit IgG (H+L), coupled to Alexa 568 | 1:500 | Invitrogen, Ref. A-10042 |
|  | Donkey anti-rabbit IgG (H+L), coupled to Cy5 Affinity Pure | 1:500 | Jackson ImmunoResearch, Ref. 711-175-152 |

### Double Immunofluorescence

After tissue permeabilization and blocking of unspecific binding, the sections were incubated with a cocktail of the primary antibodies, chicken anti-GFP and either rabbit anti-Foxg1 or rabbit anti-Nkx2.1 (Table 1), diluted in PBS-Tx for 72 hours at 4°C, under gentle agitation.

Following this, sections were washed and then incubated in a cocktail of fluorescence secondary antibodies, goat anti-chicken couple to Alexa 488 and donkey anti-rabbit coupled to Alexa 568 (Table 2), diluted in PBS-Tx for 90 minutes at room temperature. Finally, the sections were rinsed and mounted as explained above, and finally coverslipped using an antifading mounting medium (Vetashield Hardset Antifade mounting medium, Vector Laboratories Ltd.).

### Primary antibody characterization

See Table 1 for a list of all primary antibodies employed. All antibodies were validated on Western blots by the respective manufacturer, and produced specific staining patterns identical to those observed using in situ hybridization.

The chicken anti-GFP antibody recognized a single band of 25 KDa on Western blots of HEK293 transfected cell lysates, and a band at the same molecular weight on Western blots of transgenic mouse spinal cords (manufacturer’s data sheet). No staining was seen in ‘naïve’ non-transfected cells. This antibody has been successfully used to detect enhanced GFP in Viaat-eGFP knockin transgenic mice (Aresh et al., 2017). In our material from Otp-eGFP knockin mice, it produces a pattern identical to that of Otp by both in situ hybridization and immunohistochemistry (Fig. 2).

**Figure 2.**
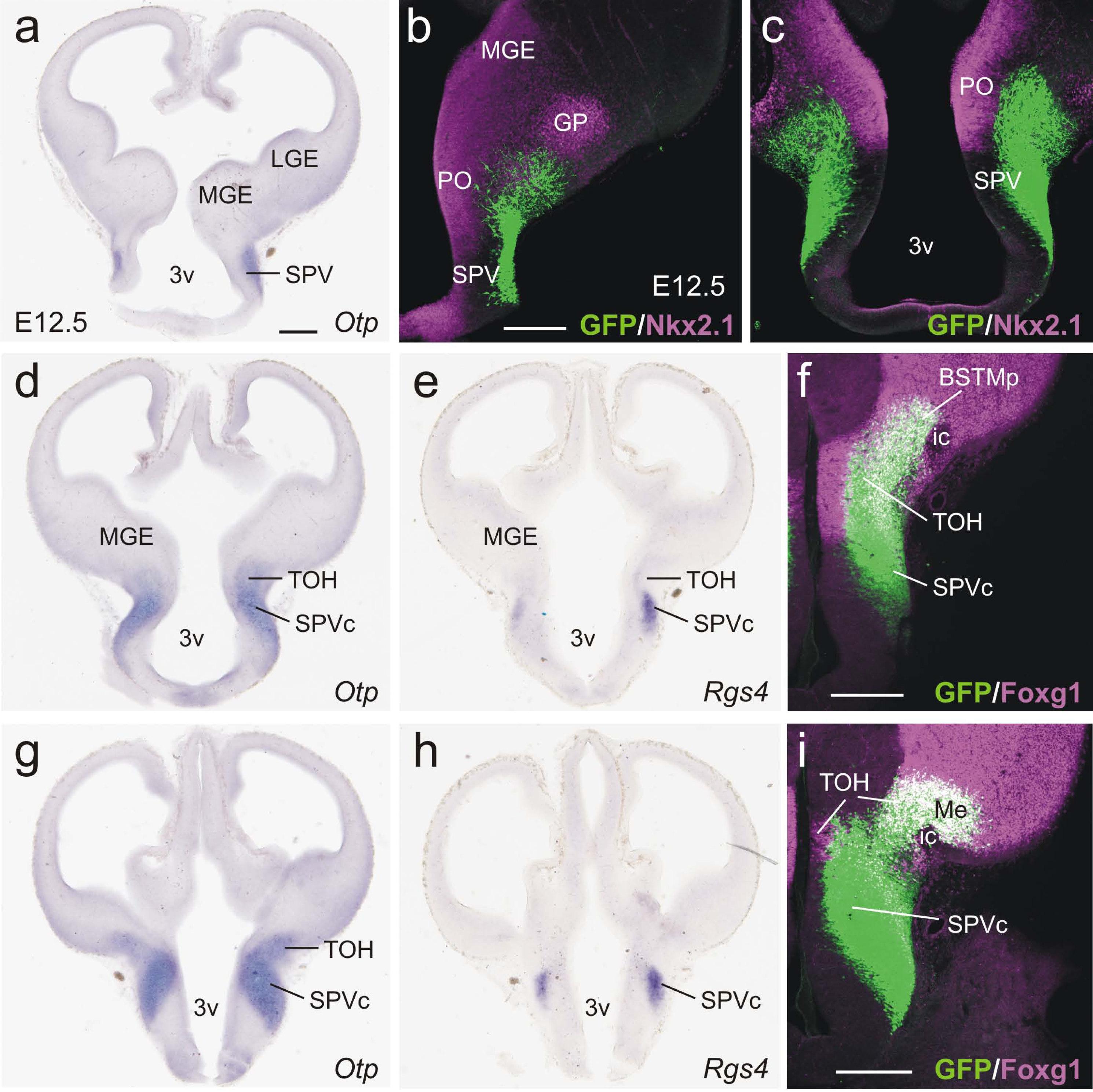
**a**,**d**,**e**,**g**,**h:** Images of frontal sections of wild type mice at E12.5 hybridized for either Otp or Rgs4. **b**,**c:** Confocal images of frontal sections of Otp-eGFP embryos at E12.5, double labeled for GFP (green) and Nkx2.1 (magenta) (double immunofluorescence). The GFP labeling in these transgenic embryos highly resembles that observed with in situ hybridization for Otp in wild type animals. In both cases, high expression is observed in SPV, but some cells are also observed at the surface of the subpallium (these are more clearly visualized in the transgenic embryos). **f**,**i:** Confocal images of frontal sections of Otp-eGFP embryos at E12.5, double labeled for GFP (green) and Foxg1 (magenta). Note the domain rich in Foxg1 (ventricular zone plus mantle) and GFP (mantle), here defined as TOH, which extends into part of the extended amygdala (posterior BSTM and Me). In contrast to TOH, the SPV core only expresses GFP. See text for more details. For abbreviations see list. Scales: bar in a = 200 µm (applies to a, d, e, g, h); bar in b = 100 µm (applies to b, c); bar in f, I = 100 µm.

The rabbit anti-GFP antibody was tested by the manufactured using immunoelectrophoresis resulting in a single precipitin arc against anti-Rabbit Serum and purified and partially purified Green Fluorescent Protein (*Aequorea Victoria*).

The rabbit anti-Foxg1 antibody recognizes a band of 50 KDa on Western blots of mouse brain tissue lysate, which is blocked with the addition of the immunizing peptide (manufacturer’s data sheet). In mouse brain sections, it produces a staining pattern identical to that observed with in situ hybridization for Foxg1 (Dou et al., 1999; Crossley et al., 2001).

The rabbit anti-TTF1 (anti-Nkx2.1) antibody recognizes a single band of about 42 kDa on Western blots of rat brain samples. The staining pattern observed in the developing mouse forebrain sections with this antibody is identical to that obtained with in situ hybridization (García-López et al., 2008).

The rabbit anti-Otp antibody recognized a band of 34 KDa on Western blots of Jurkat cell lysates (manufacturer’s data sheet). The staining pattern in developing mouse brain sections with this antibody is identical to that obtained by in situ hybridization (Fig. 2).

The mouse anti-Islet1 antibody was raised against the C-terminal residues 178–349 of rat Islet-1, produces a comparable staining pattern in rat and chicken brain, and recognizes both Islet1 and Islet2 (Thor et al., 1991; Varela-Echavarría et al., 1996). The staining pattern obtained with this antibody in the developing mouse forebrain (Elshatory and Gan, 2008; Bupesh et al., 2011b) is identical to that obtained with in situ hybridization (Stenman et al., 2003).

The mouse anti-RC2 antibody recognizes the intermediate filament nestin (radial glial marker), and has been tested on Western blot of mouse tissue, where it recognizes a band of about 295 KDa (manufacturer’s data sheet).

### In situ hybridization: conventional method

Brain sections were processed for in situ hybridization using digoxigenin-labeled riboprobes and following a procedure previously described (Abellán et al., 2014). The antisense digoxigenin-labeled riboprobes were synthesized using Roche Diagnostics (Germany) protocols from cDNAs of different genes of interest, including Otp, Sim1 and Rgs4 (Table 3).

**Table 3.**
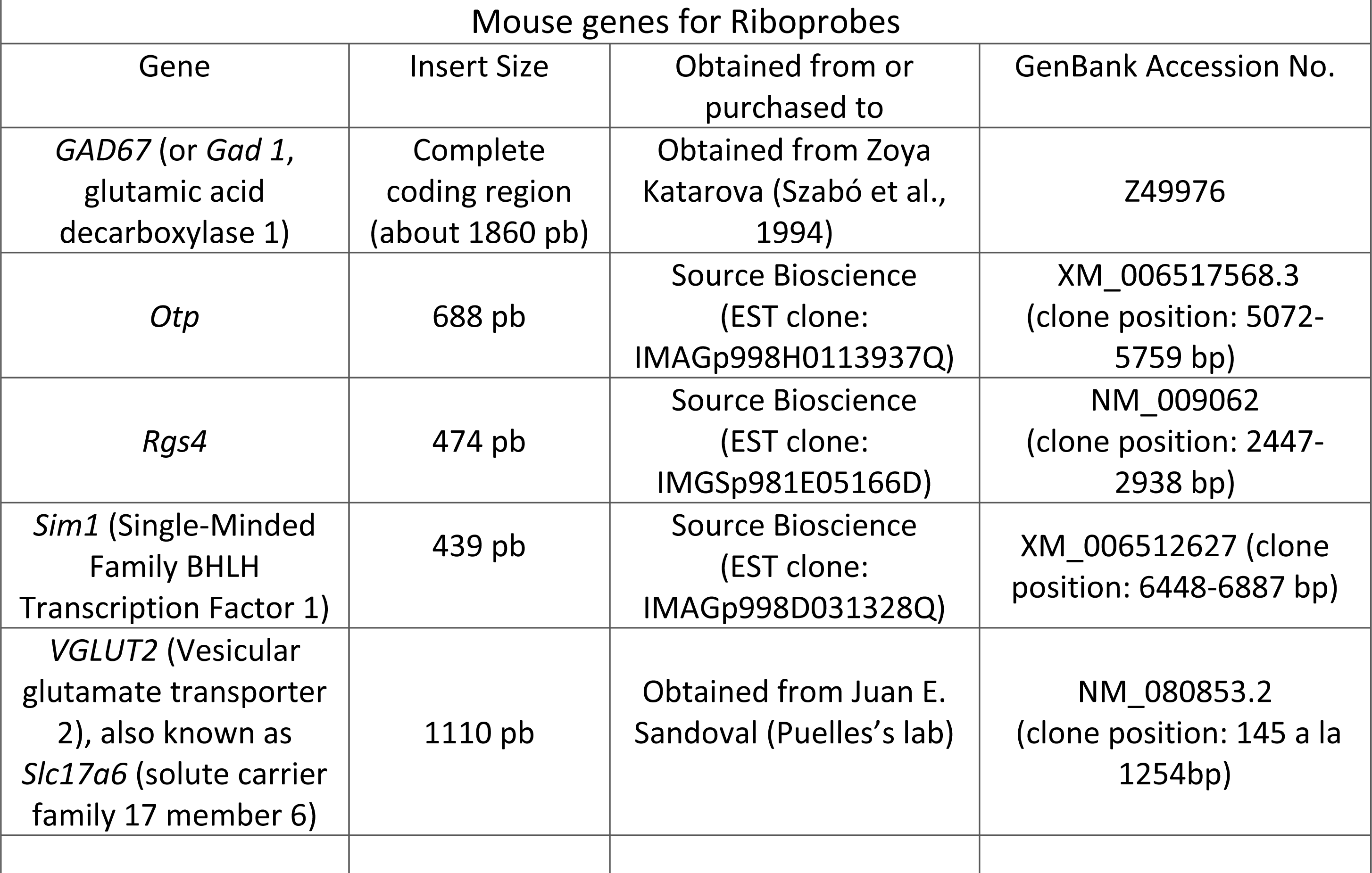
Genes for riboprobes.

Before starting the hybridization, the tissue was permeabilized with 0.1% Tween-20 in PBS (PBT; 0.1M), washing 3 times for 10 minutes each. Following this, the sections were prehybridized in hybridization buffer (HB), containing 50% deionized formamide (Sigma-Aldrich), 6.5% standard saline citrate (pH=5; 2M), 1% ethylenediaminetetraacetic acid (EDTA; pH=8; 5mM; Sigma-Aldrich), 0.2% Tween-20, 1mg/mL of yeast tRNA (Sigma-Aldrich), 100 µg/mL of heparin (Sigma-Aldrich), completed with RNase and DNase water (Sigma-Aldrich), for 2-4 hours at 58°C. Then, the sections were hybridized overnight at 58°C in HB containing 0.5-1 µg/mL of the corresponding riboprobe. The next day, the hybridized sections went through a series of washes: first in HB at 58°C, then in a mix 1:1 of HB and MABT (1.2% Maleic acid, 0.8% NaOH, 0.84% NaCl, and 0.1% Tween-20) at 58°C and finally in MABT at room temperature.

Then, the sections were blocked to avoid unspecific binding using a solution containing blocking reagent and sheep serum in MABT, for 4 hours at room temperature. Following this, the sections were incubated, overnight at 4°C, with a sheep anti-digoxigenin antibody conjugated with alkaline phosphatase (Roche Diagnostics) diluted 1:3500 in MAB). After washing in the same buffer, the sections were finally revealed with BM purple (Roche Diagnostics) and then mounted on glycerol gelatin mounting medium (Sigma-Aldrich).

### Double and triple labeling: Single and Double Immunofluorescence combined with indirect FISH

Some of the sections processed for double or triple labeling were first processed for fluorescence in situ hybridization (FISH), using an indirect method. In particular, we did this to detect Sim 1, GAD67 (glutamic acid decarboxylase 67, to detect GABAergic neurons) or VGLUT2 (vesicular glutamate transporter 2, to detect glutamatergic cells) (see Table 3 for details). For this procedure, we used antisense digoxigenin-labeled riboprobes, which were synthesized as explained above.

Before starting the hybridization, the tissue was permeabilized with 0.1% Tween-20 in PBS (PBT; 0.1M), by washing 3 times for 10 minutes each. Then, the sections were pre-hybridized in HB for 2-4 hours at 58°C. Then, the sections were hybridized overnight at 58°C in HB containing 1 µg/mL of the corresponding riboprobe. The next day, the sections were washed with HB for 30 minutes at 58°C. After that, we continued washing using saline sodium-citrate buffer (SSC; pH = 7.5, 0.02M) 3 times for 15-20 minutes at 58°C, followed by 1 wash in the same buffer for 15 minutes at room temperature and 1 wash with Tris buffer (TB, 0,05M, pH 7.6) for 15 minutes at room temperature. The activity of the endogenous peroxidase was inhibited as described above but diluting the hydrogen peroxide in TB. After, the sections were washed with Tris-NaCl-Tween buffer (TNT; 10% TB, pH 8, 0.1M; 0.9% NaCl; 0.05% Tween-20) for 15 minutes at room temperature. Following this, the sections were incubated in a blocking solution consisting of 20% blocking reagent (BBR) and 20% of sheep serum in TNT (TNB) for 2-4 hours, and with a sheep anti-digoxigenin antibody (Roche Diagnostics, Basel, Switzerland) conjugated to the peroxidase enzyme, diluted 1:200 in TNB overnight at 4°C and under gentle agitation. After washing with TNT 3 times for 10 minutes, the slices were incubated with TSA Working solution, containing tyramide conjugated to Cy3 (diluted 1:20; TSA Plus Cy3 System, Perkin Elmer, United States) for 10 minutes. Finally, the sections were rinsed and processed for single or double immunofluorescence, as described next.

After the fluorescent in situ hybridization, the sections hybridized for GAD67, VGLUT2 or Sim1 were processed for single or double immunofluorescence, in order to detect GFP (double labeling) and Foxg1 (triple labeling). The procedure was similar to that described above (section on double immunofluorescence). The primary antibodies were chicken anti-GFP and rabbit anti-Foxg1 (Table 1), while as secondary antibodies we employed goat anti-chicken coupled to Alexa 488, and donkey anti-rabbit couple to Cy5 (Table 2).

### Digital photographs and figures

Digital microphotographs from immunohistochemical sections were taken on a Leica microscope (DMR HC, Leica Microsystems GmbH) equipped with a Zeiss Axiovision Digital Camera (Carl Zeiss, Germany). Low magnification digital microphotographs were obtained with a stereoscopic microscope (2000C Zeiss) equipped with a Canon Digital Camera (EOS 450D, Canon Inc., Japan). Serial images from fluorescent material were taken with a confocal microscope (Olympus FV1000, Olympus Corporation, Japan). Selected digital immunohistochemical images were adjusted for brightness and contrast with Corel PHOTO-PAINT 2012 or 2019 (Corel Corporation, Canada), while the fluorescent ones were adjusted and extracted using Olympus FV10-ASW 4.2 Viewer (Olympus Corporation). Finally, the figures were mounted using CorelDraw 2012 or 2019 (Corel Corporation). The schemes (or drawings) included in the figures were made by means of Power Point (Microsoft Corporation, United States of America), and were based on selected immunostained microphotographs of representative brain levels on frontal, sagittal or horizontal planes.

### Nomenclature

For embryonic domains and axis, we follow the prosomeric model (Puelles and Rubenstein, 2015) as well as our previous studies on amygdalar development in mouse (García-López et al., 2008; Bupesh et al., 2011a,b). This implies using the terms ‘dorsal’ and ‘ventral’ following topological coordinates (Nieuwenhuys and Puelles, 2016), which in the forebrain are about 90 degrees shifted with respect to the classical topographic terms. For example, the dorsal part of the paraventricular hypothalamic nucleus as used here corresponds to the anterior part of this nucleus in classical nomenclature. We did this when referring to general regions and divisions, but for specific nuclei and areas within them, we employed the terminology of the Franklin and Paxinos’ mouse brain atlas.

## Results

### The TOH: a new embryonic domain at the frontier between telencephalon and hypothalamus

The results in zebrafish on a new embryonic domain (called ‘optic recess region’) at the transition between telencephalon and hypothalamus, expressing both Otp and Foxg1, prompted us to investigate whether a similar division exists in mouse. To that aim, we processed brain sections from Otp-eGFP mice at different embryonic stages for double immunofluorescence to detect GFP (expressed in Otp-lineage cells) and the transcription factor Foxg1. To better understand the location of the GFP expression domains, we also compared the expression of GFP/Otp with other region-specific transcription factors such as Nkx2.1, Islet1, and Sim1 (**Figs. 1-3**).

At early stages (E12.5), GFP labeling in the forebrain of the transgenic mice recapitulated the mRNA expression of Otp, being highly expressed in a forebrain domain previously identified as the supraopto-paraventricular division (SPV) or simply paraventricular division of the alar hypothalamus (**Figs. 1-3**). This GFP/Otp expressing domain was surrounded by other domains expressing Nkx2.1 and/or Islet1 (**Figs. 1, 3**), as follows: (1) Dorsally, SPV was bounded by the preoptic region of the telencephalon, characterized by expression of Nkx2.1 and Islet1 (**Figs. 1b, 3**). (2) Ventrally, it was bordered by the subparaventricular region and the basal hypothalamus, characterized by expression of either Islet1 or Nkx2.1, respectively (**Figs. 1b, 3**). (3) Caudally, it was bounded by the prethalamus, characterized by expression of Islet1 (**Fig. 3**). However, double labeling of GFP (Otp) and Foxg1 showed a clear overlapping of both in a dorsal subdivision of classical SPV (**Fig. 1a**,**c**). As seen in sagittal sections, the GFP/Foxg1 overlapping area was located at the transition between the telencephalon and hypothalamus, covering both peduncular and prepeduncular (terminal) prosomeric subdivisions of the secondary prosencephalon (**Fig. 1a**,**c;** for reference, prosomeric divisions are labeled, at hypothalamic levels, as PHy and THy in **Fig. 1d**). In frontal sections, we confirmed that the vast majority of the cells in the mantle of this transition region truly coexpress GFP and Foxg1 (**Fig. 4**). This mouse forebrain division corresponds to the ‘optic recess region’ of zebrafish. Since it produces not only part of the eye vesicle and pedicle (**Fig. 1a**), but also part of the telencephalon and hypothalamus (as explained later), we called this new division in mouse the ‘telencephalo-opto-hypothalamic domain’ or TOH (**Fig. 1a**,**c**,**f**). Like the corresponding division in the zebrafish, the mouse TOH showed not only overlap of GFP/Otp with Foxg1, but also with Sim1 (**Fig. 3**). Overall, this new division was distinct from the subpallial telencephalon because of its expression of Otp and Sim1, but lack of expression of Nkx2.1 and Islet1 (**Figs. 1-5**).

**Figure 3.**
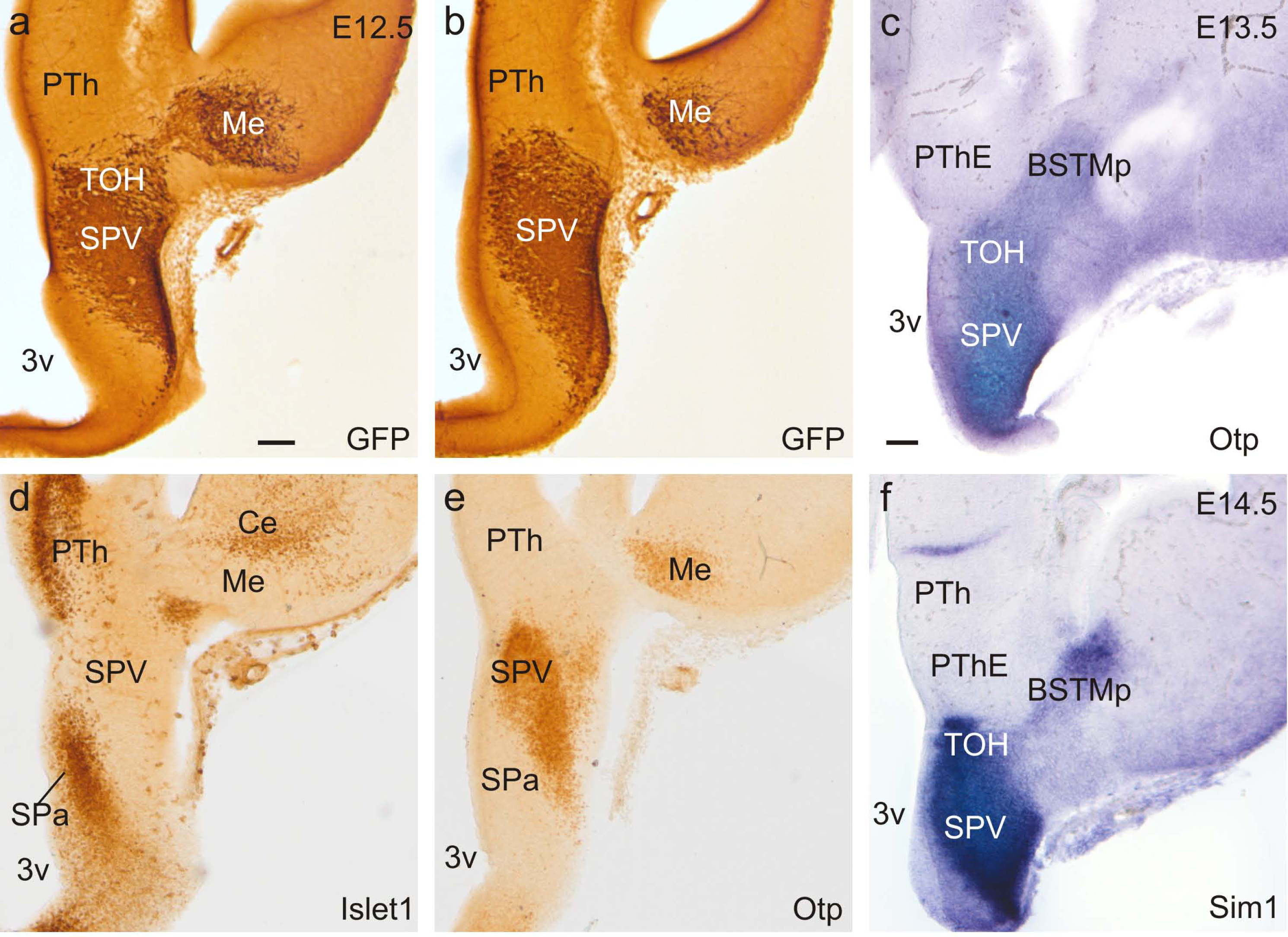
**a**,**b:** Frontal sections of Otp-eGFP embryos at E12.5, stained by way of immunohistochemistry for GFP. **d**,**e:** Frontal sections of wild type embryos at E12.5, stained by way of immunohistochemistry for Islet1. Comparison between these sections shows that the GFP-rich SPV area (including the TOH subdivision) is bounded by areas rich in Islet1. GFP and Islet1 also occupy largely non-overlapping areas in the amygdala. **c**,**f:** Frontal sections of wild type embryos at E13.5 (c) or E14.5 (f), hybridized for Otp or Sim1. See text for more details. For abbreviations see list. Scale bar in a = 100 µm (applies to a, b, d, e); bar in c = 100 µm (applies to c, f).

**Figure 4.**
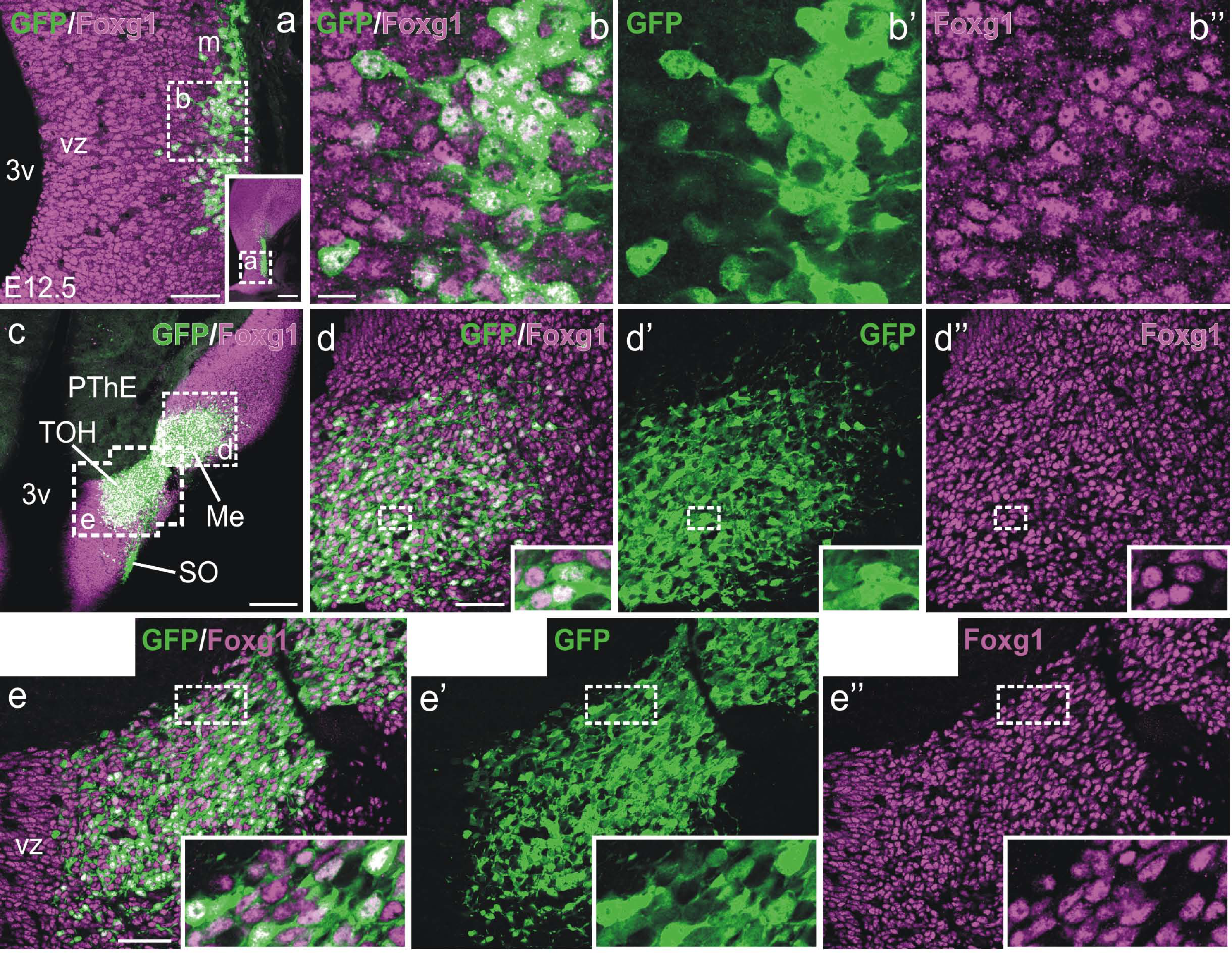
Confocal images of frontal sections of Otp-eGFP embryos at E12.5, double labeled for GFP (green) and Foxg1 (magenta) (double immunofluorescence), at the level of TOH (including part of Me). Squared areas in a and c are shown at higher magnification in b-b’’, d-d’’ and e-e’’. Insets in d-d’’ and e-e’’ show additional details of cells (e-e’’ show a photomontage). These details show that the vast majority of GFP cells co-express Foxg1. For abbreviations see list. Scales: a = 50 µm; insert in a (panoramic view) = 200 µm; b = 10 µm (applies to b-b’’); c = 200 µm; d, e = 50 µm (applies to d-d’’ and e-e’’).

Moreover, this division was different from the hypothalamus because of its high expression of Foxg1 in both ventricular zone (vz) and mantle (**Figs. 1, 2, 4**). Thus, our observations implied a redefinition of the boundary between telencephalon and hypothalamus, and let us to conclude that the classical SPV (represented in **Fig. 1e**) is divided into two sectors (**Fig. 1f**): (1) a dorsal subdivision or TOH (non-hypothalamic) and (2) a ventral subdivision, formed by the main or core part of SPV (hypothalamic). Both sectors were also distinguished based on their distinct expression of Rgs4 (a regulator of G protein signaling 4, previously considered to be distinctly expressed in the peduncular prosomeric part of classical SPV): our data showed that Rgs4 is highly expressed in the mantle of the core part of SPV (at peduncular prosomeric levels), but it is very weak in TOH (**Fig. 2e**).

To better understand the radial extension of TOH, we compared Otp/GFP with radial glial fiber disposition (using a RC2 antibody) in different section planes through the forebrain. Our results showed that horizontal sections were particularly useful to visualize the TOH in its whole radial extension, from the ventricle to the subpial surface (**Figs. 6, 7**). Comparison of Otp with RC2 in this same plane at E12.5 demonstrated that the TOH radial histogenetic division extends from its vz, covering the third ventricle and adjacent interventricular foramen, to the surface in the ventral and caudolateral telencephalon, an area that includes at least part of the medial amygdala complex (**Fig. 6**). Based on analysis of sagittal, frontal and horizontal sections at E12.5 and later stages (**Figs. 1, 2, 4, 6-8**), the posterior BSTM, the anterior medial amygdala (MeA) and at least the medialmost part of the posterior medial amygdala (MeP) are located within the same radial domain in TOH (**Fig. 6**). In addition, the use of different section planes allowed the distinction between the peduncular and pre-peduncular (or terminal) prosomeric parts of TOH, since these are best appreciated in sagittal sections (labeled as reference in the hypothalamus [PHy, THy] in **Fig. 1d**). Regarding the peduncular prosomeric part, it appears to produce the dorsal paraventricular nucleus (adjacent to the third ventricle; **Fig. 1c**), the posterior BSTM (adjacent to the interventricular foramen; **Figs. 1a**,**b; 2f; 6a**,**b; 7a**,**b; 8b**), and part of the medial amygdala (laterally located) (**Figs. 2i; 4c; 7a-c; 8a**). With respect to the pre-peduncular (terminal) prosomeric part of TOH, it produces a ventral part of the preoptic region (POv, **Figs. 1c; 8b**).

**Figure 5.**
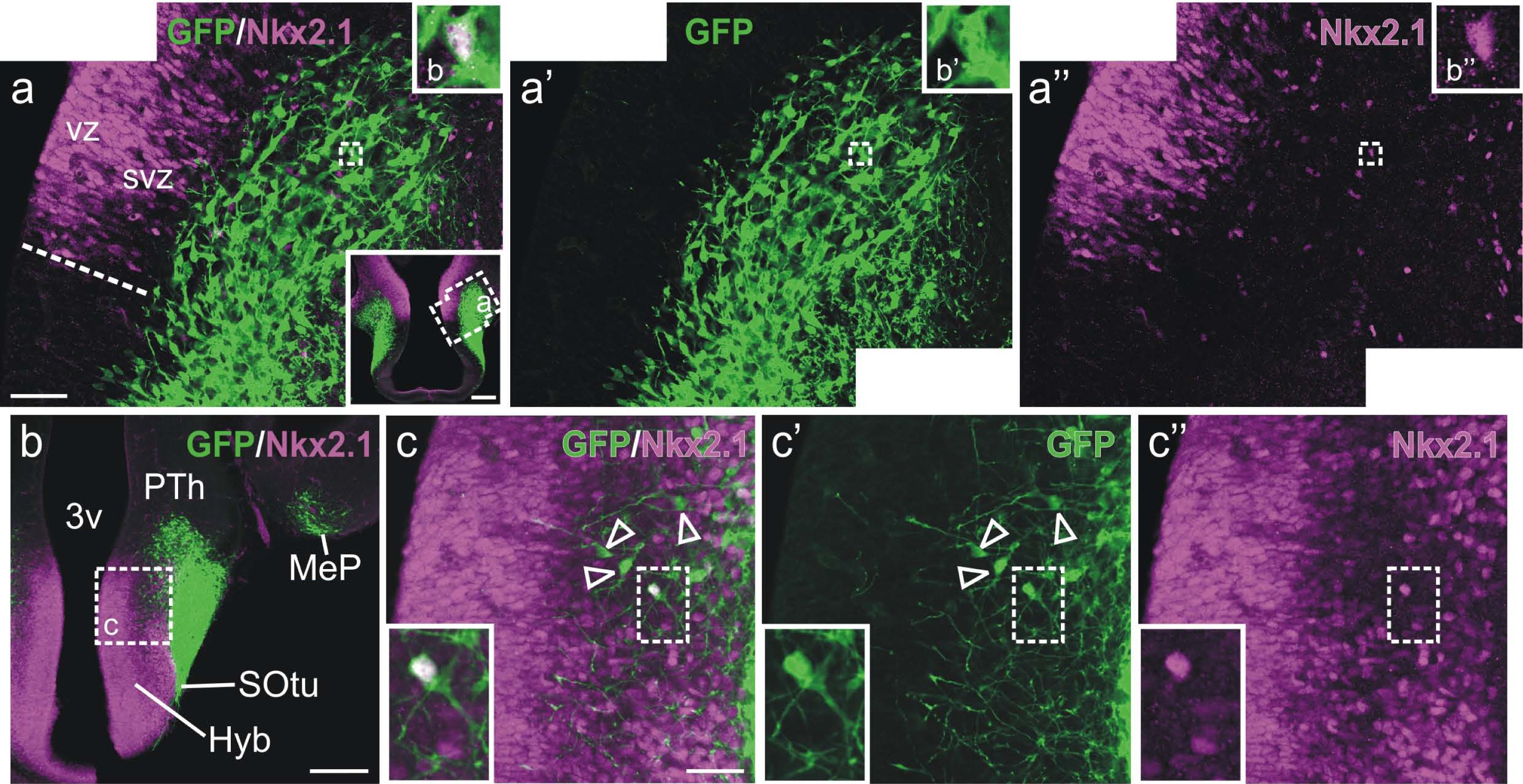
Confocal images of frontal sections of Otp-eGFP embryos at E12.5, double labeled for GFP (green) and Nkx2.1 (magenta) (using double immunofluorescence), at the level of the telencephalic subpallium (a-a’’) or the basal hypothalamus (b,c-c″). The pallido-preoptic subdivisions of the subpallium and the basal hypothalamus show their typical expression of Nkx2.1 in ventricular zone and mantle. In both regions, the majority of GFP cells locate in the mantle and do not co-express Nkx2.1, with a few exceptions shown in the insets. For abbreviations see list. Scales: bar in a = 50 µm (applies to a-a’’); insert in a (panoramic view) = 200 µm; b = 200 µm; c = 20 µm (applies to c-c’’).

**Figure 6.**
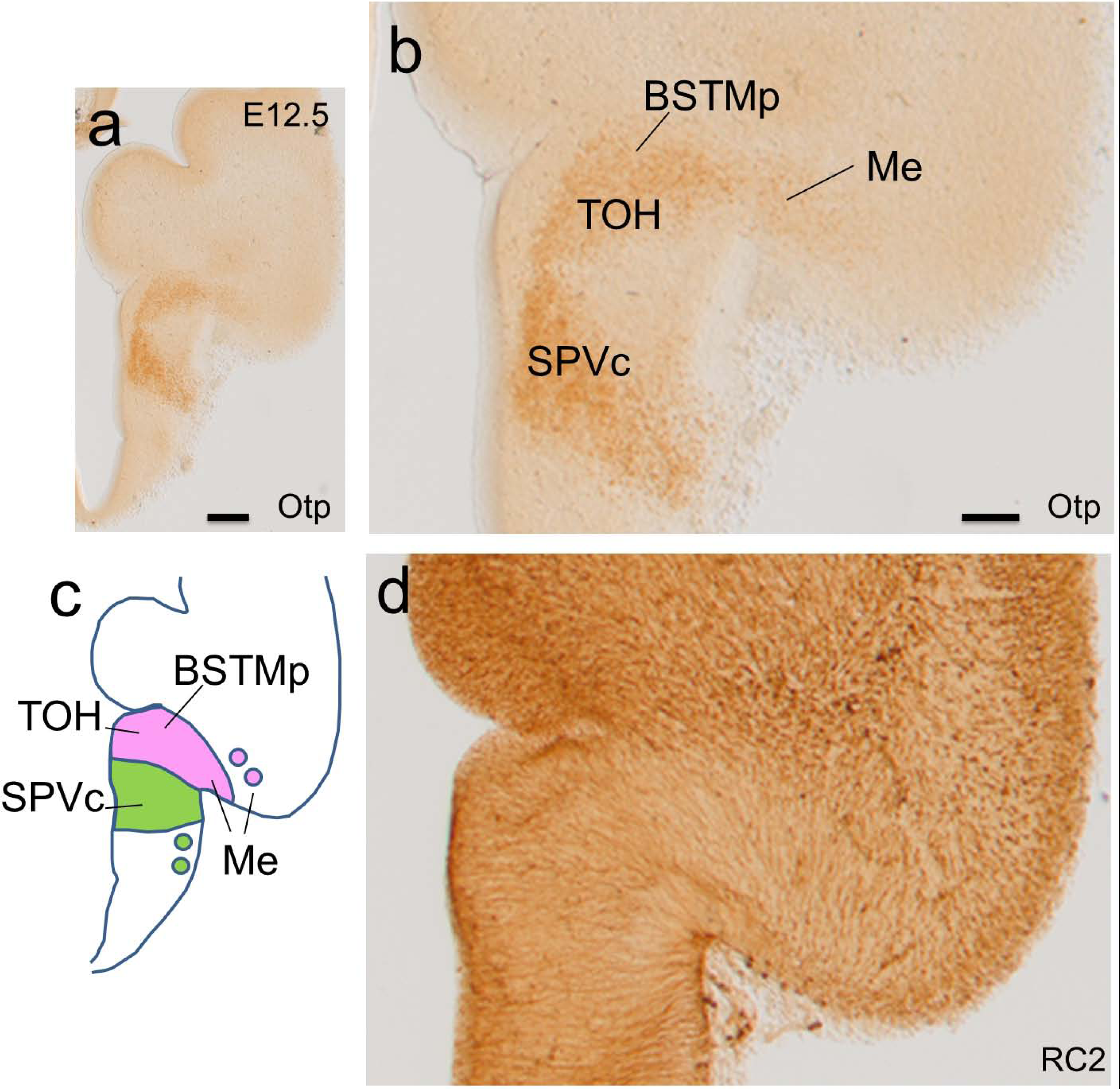
**a**,**b**,**d:** Horizontal adjacent sections of a wild type embryo at E12.5, processed for immunohistochemistry to detect Otp or RC2 (a radial glial marker). Two expression domains of Otp are observed, one in TOH and another one in SPV core. Comparison with radial glial fibers show that TOH extends radially into a caudoventral part of the telencephalon, that encompasses part of the medial extended amygdala. The TOH radial histogenetic domain is represented in a scheme in c. See text for more details. For abbreviations see list. Scales: a = 200 µm; b = 100 µm (applies to b, c).

**Figure 7.**
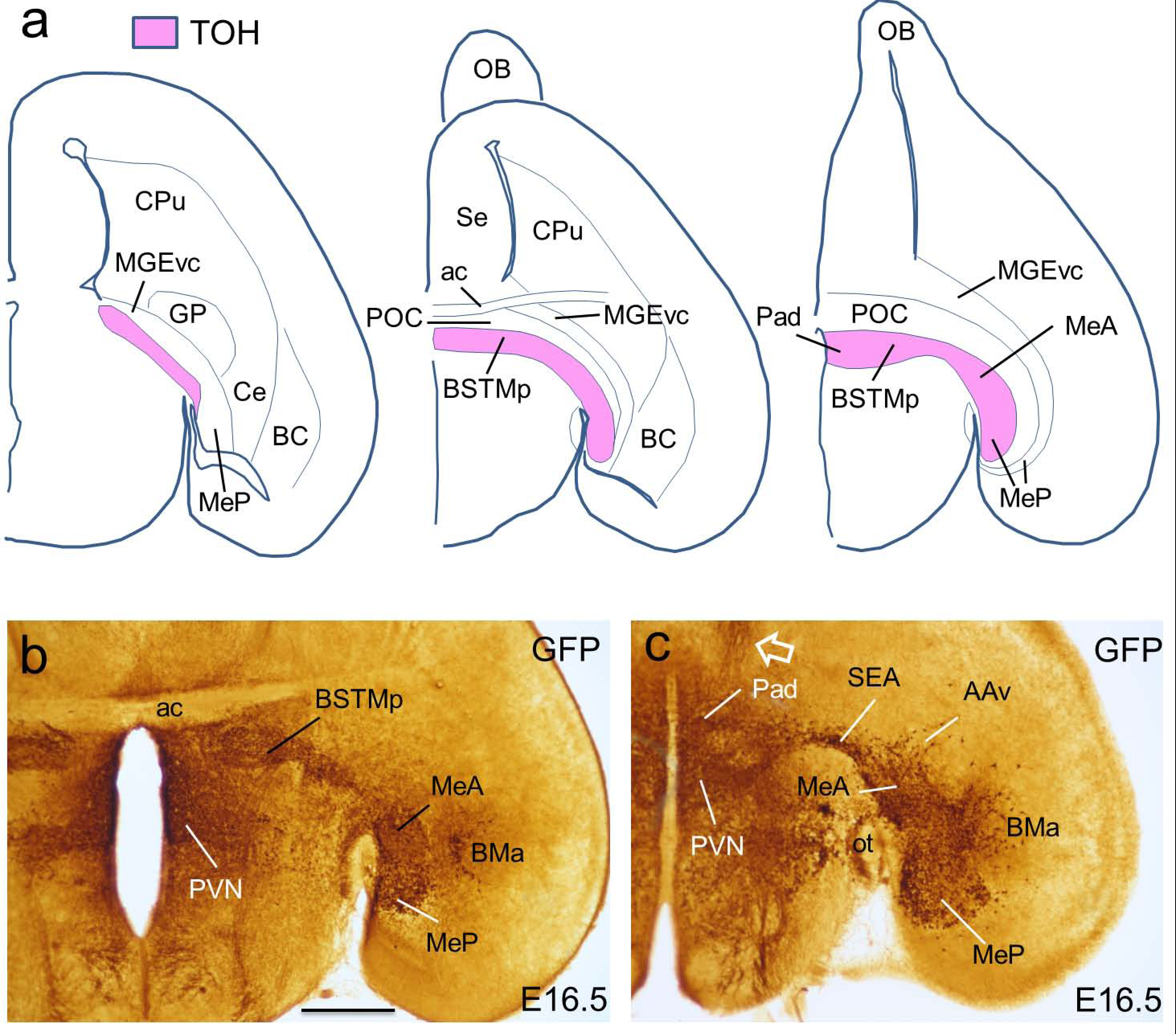
**a:** Schemes of horizontal sections of the embryonic brain at E16.5, representing the TOH radial histogenetic domain. The central section is at the level of the anterior commissure, that on the left is above, while that of the right is below the anterior commissure. In these sections, rostral is to the top. These schemes are based on horizontal sections from Otp-eGFP embryos stained for GFP (using immunohistochemistry), from which we selected those in **b** and **c**. These sections allow a better comprehension of the TOH division, which includes areas previously considered part of the hypothalamus or the telencephalon (including part of the medial extended amygdala). The arrow in c point to a labeled fiber tract projecting from TOH to the septum. See text for more details. For abbreviations see list. Scale: b = 500 µm (applies to b, c).

In addition, our analysis of sagittal and frontal doubled-labeled sections through the forebrain, from E12.5 on, allowed us to suggest that TOH is a major source of double-labeled GFP/Foxg1 tangentially migrating cells, part of which enters into the telencephalic subpallium (located dorsal to TOH), while another part appears to move ventrally, into the hypothalamus (**Figs. 1, 2, 5**). Regarding those that appeared to enter the subpallium, at early stages these cells were already seen in the pallidopreoptic region, characterized by expression of Nkx2.1 (**Figs. 2, 5)**.

However, GFP cells were primarily located superficially to Nkx2.1-expressing progenitors (located at the ventricular and subventricular zones) of the preoptic and ventrocaudal pallidal (diagonal) embryonic divisions (**Figs. 2b**,**c, 5a**; as a reference, the latter division locates ventral to incipient globus pallidus and produces a part of the medial extended amygdala). Moreover, many GFP cells displayed a migratory neuroblast morphology and showed a clear tangential disposition, with a leading process directed dorsally (**Fig. 5a**). The vast majority of GFP cells in the telencephalon were different from Nkx2.1 expressing cells (**Fig. 5a**) and instead expressed Foxg1 (**Fig. 4**; see further details in next section). Subsets of these cells reached the subpallial part of the preoptic area, subpallial parts of the extended amygdala, and pallial parts of the amygdala, such as the cortical areas and basomedial nucleus (**Figs. 7, 8**). Nevertheless, we also observed extremely few cases of GFP cells in the subpallium co-expressing Nkx2.1 (**Fig. 5b’**).

**Figure 8.**
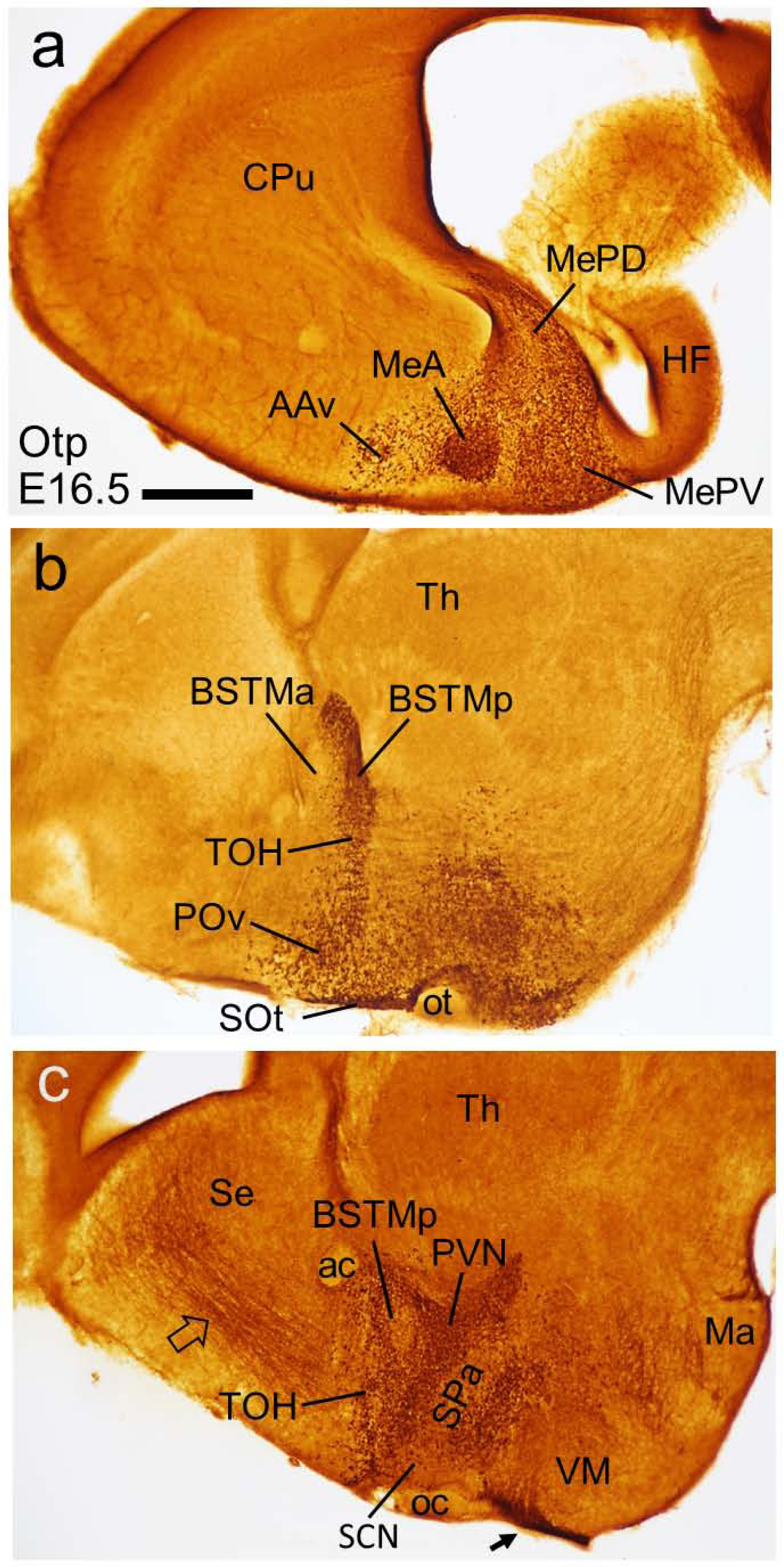
Sagittal sections from Otp-eGFP embryos at E16.5, stained for GFP using immunohistochemistry, from lateral (top) to medial (bottom) levels. In these sections, rostral is to the left and dorsal to the top. The TOH includes a ventral preoptic subdivision, a posterior subdivision of BSTM and, more laterally, a part of the medial amygdala. At more medial levels (not shown here, but seen in Fig. 15), it also includes a dorsal subdivision of the paraventricular nucleus. In contrast, the main part of this nucleus (PVN) and the supraoptic nucleus belongs to SPV core. The arrow in c points to a labeled fiber tract projecting to the septum. See text for more details. For abbreviations see list. Scale: a = 500 µm (applies to a-c).

Furthermore, although the vast majority of the GFP cells of the telencephalon and TOH were double-labeled for Foxg1, we could also observe a few GFP cells in a subpial position, which do not co-express Foxg1 (**Figs. 9, 10b**). We interpret that GFP cells (Foxg1-negative) of these areas are derived from the SPV core, as discussed later.

**Figure 9.**
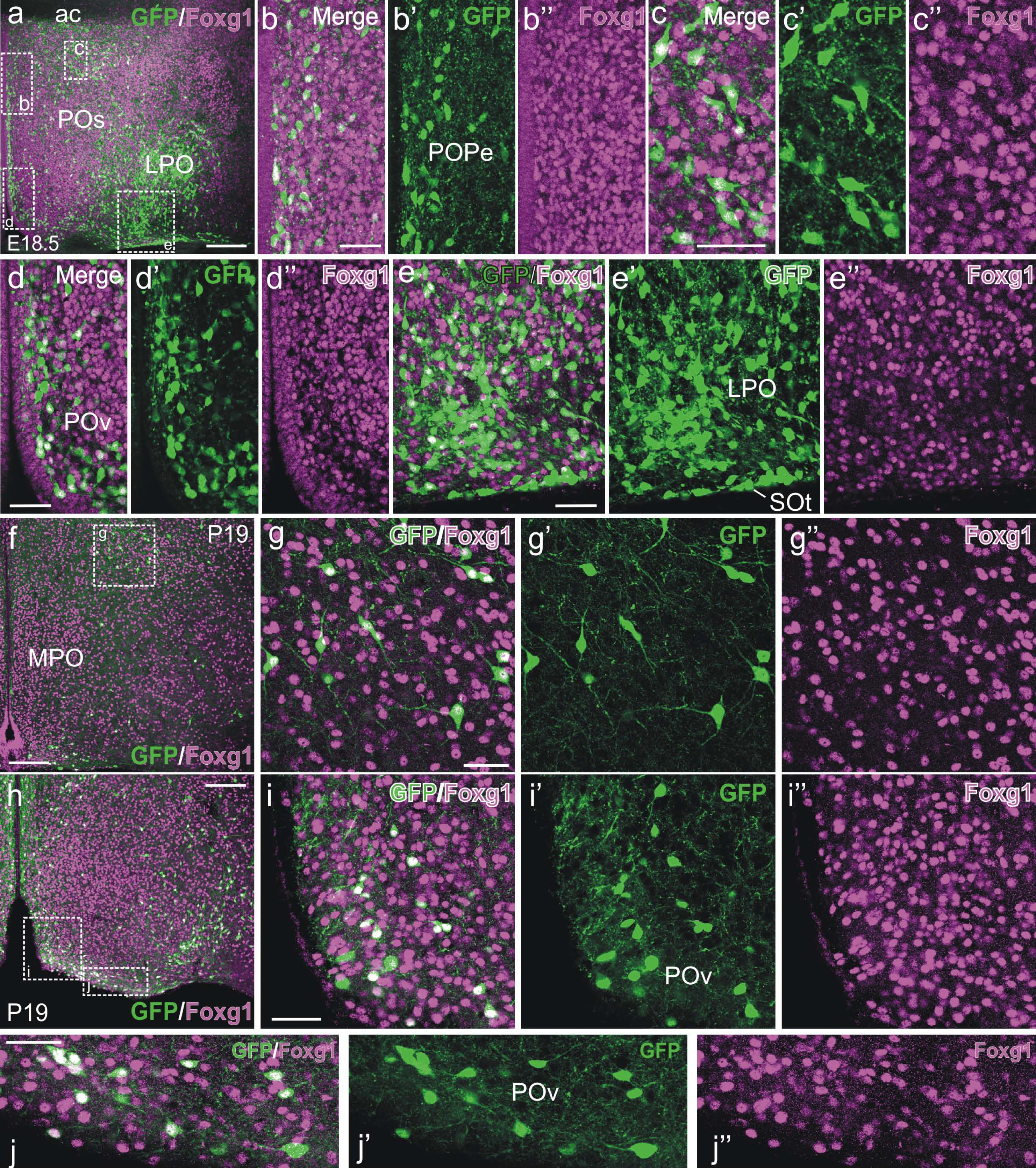
Confocal images of frontal sections of E18.5 and P19 Otp-eGFP mice, at the level of the preoptic region, double labeled for GFP (green) and Foxg1 (magenta). Panoramic views are shown in a, f and h, while details of the squared areas in these are shown in b-b’’, c-c’’, d-d’’, e-e’’, g-g’’, i-I’’ and j-j’’. Double labeled cells tend to accumulate in the ventral preoptic area, which is part of TOH, although some cells are also seen in dorsal parts of the preoptic area. Scattered single GFP-labeled cells are also seen in dorsal and ventral preoptic areas. The terminal part of the supraoptic nucleus, containing GFP single labeled cells, is seen at the surface in e,e’. See text for more details. For abbreviations see list. Scales: a = 200 µm (applies to a,f,h); b = 50 µm (applies to b-b’’); c = 50 µm (applies to c-c’’); d, e, g, i, j = 50 µm (applies to d-d’’, e-e’’, g-g’’, i-i’’, j-j’’).

**Figure 10.**
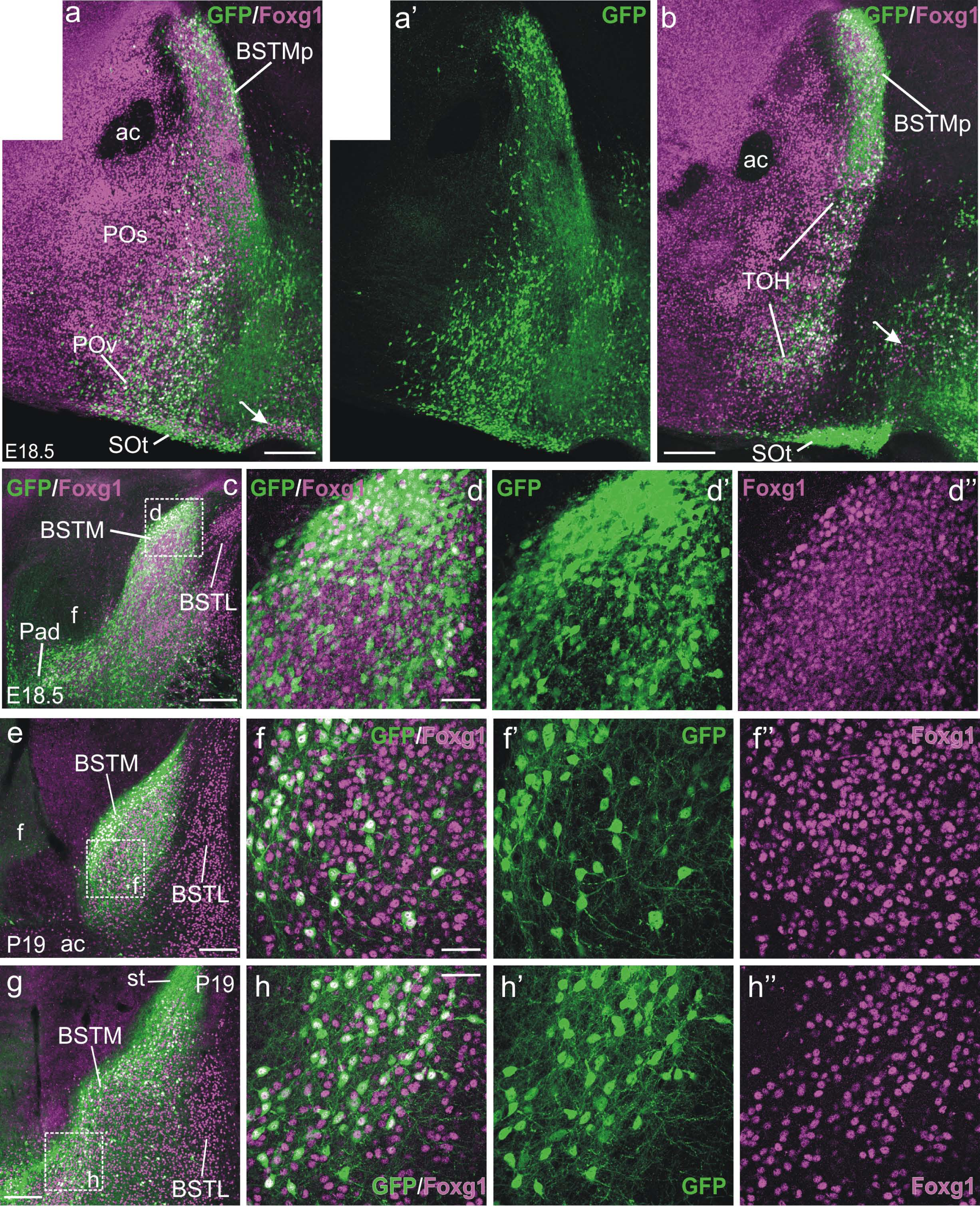
Confocal images of sagittal (a,a’,b) and frontal (c-h’’) sections of E18.5 and P19 Otp-eGFP mice, at the level of the BSTM, double labeled for GFP (green) and Foxg1 (magenta) (a, a’ and b show photomontages). Details of the areas squared in c, e and g are shown in d-d’’, f-f’’ and h-h’’. Note the high number the GFP labeled cells in the posterior BSTM, most of which co-express Foxg1. Nevertheless, GFP cells in this nucleus do not distribute homogeneously, and some subareas are rich in single Foxg1 labeled cells. Scattered GFP single labeled cells are also observed. Also note the position of the main part of the terminal supraoptic nucleus (SOt) in SPV core (poor in Foxg1). Cells of SOt are single labeled for GFP, and a few appear to spread dorsally, into TOH and beyond. The arrows in a and b point to some Foxg1 cells in the hypothalamus. See text for more details. For abbreviations see list. Scales: a = 200 µm (applies to a,a’); b,c,e,g = 200 µm; d,f,h = 50 µm (applies to d-d’’,f-f’’,h-h’’).

Regarding the GFP/Foxg1 double-labeled cells that appeared to spread from TOH into the hypothalamus, these cells were found across the paraventricular and subparaventricular domains of the alar hypothalamus, as well as in the basal hypothalamus (as a reference, the latter region is characterized by expression of Nkx2.1; **Figs. 1b-d; 5b**). However, the basal hypothalamus also contained abundant GFP-labeled/Foxg1-negative cells (**Fig. 5b**). The origin of the GFP-labeled/Foxg1 negative cells of the basal hypothalamus could be double, since we observed colocalization of GFP and Nkx2.1 in some of the cells but not in others (inset in **Fig. 5c**). In addition, we also observed a remarkable continuum of Foxg1 cells without co-expression of GFP extending from the basal telencephalon into the TOH and hypothalamus (arrow in **Fig. 1d**).

### Distribution of cells derived from TOH

To study the cell populations that derive from TOH, we analyzed the detailed distribution of cells co-expressing GFP and Foxg1, focusing on late embryonic stage (E18.5) and postnatal day 19 (P19). We found several areas with a high density of GFP/Foxg1 doubled labeled cells, including a ventral part of the preoptic area (**Figs. 9, 10a,b**), a dorsal part of the paraventricular hypothalamic nucleus (**Figs. 10b**), a posterior part of BSTM (**Figs. 10b-h; 11h-h’’**), and part of the medial amygdala (**Figs. 12; 13; 14g-g’’**). All these areas are part of TOH. Subsets of GFP/Foxg1 double-labeled cells were also observed dorsal to TOH, in the subpallial part of the preoptic region (**Fig. 10a**), and ventral to TOH, in the alar and basal parts of the hypothalamus (**Figs. 15-17**). In each of these forebrain regions, GFP/Foxg1 double-labeled cells were intermingled with Foxg1-only and/or GFP-only expressing cells, in different proportions. Due to the complexity of the distribution and coexistence of these three different cell types, we organized the description in separate sections representing major forebrain regions or subregions. We paid special attention to the medial extended amygdala, which is one of the major aims of this study.

**Figure 11.**
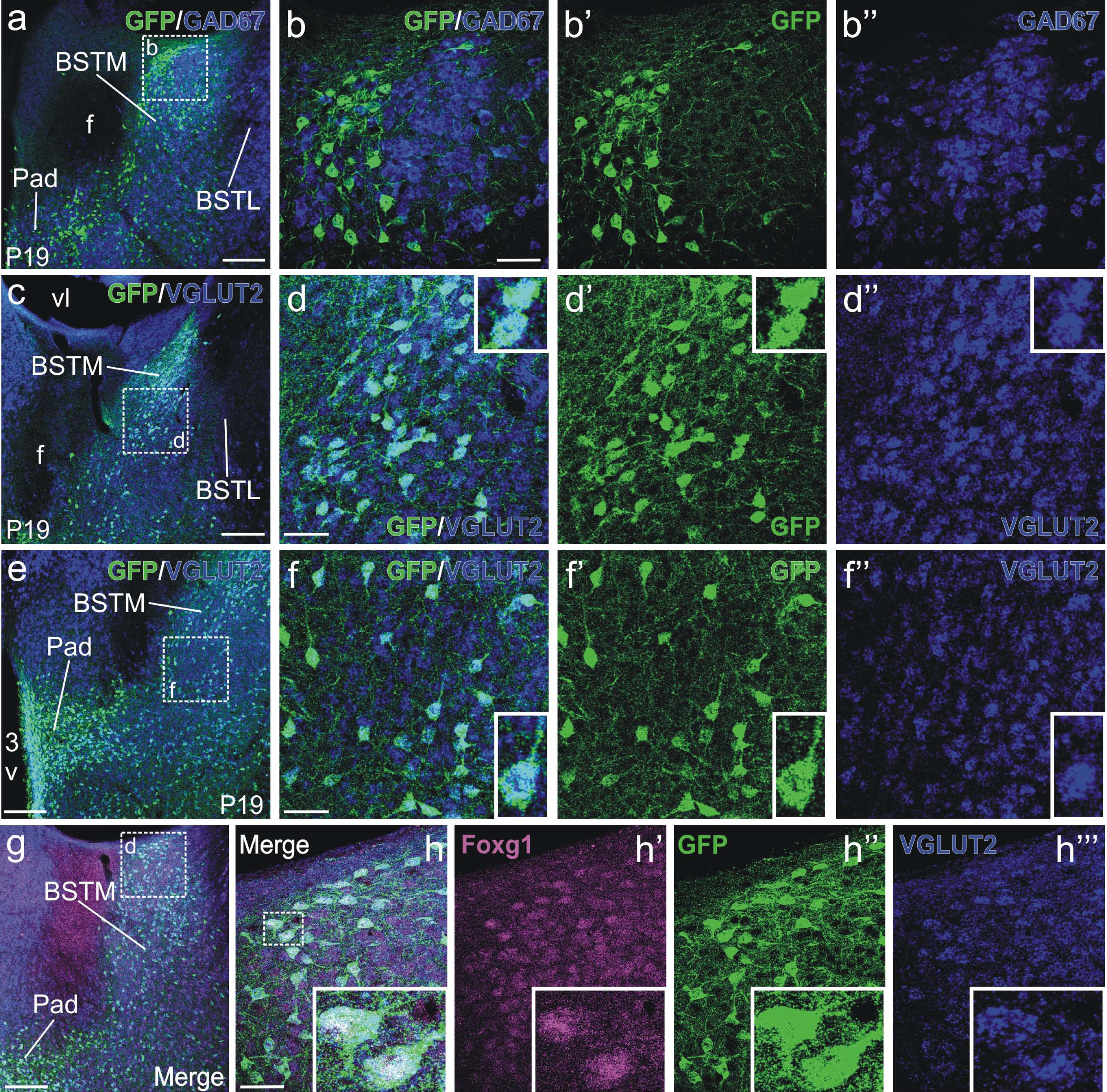
Confocal images of details of frontal brain sections of P19 Otp-eGFP mice, at the level of the BSTM, double labeled for GFP (green immunofluorescence) and either for GAD67 or VGLUT2 (blue fluorescent in situ hybridization; a-b’’ GAD67; c-f’’ VGLUT2). Triple labeling for GFP (green), Foxg1 (magenta) and VGLUT2 (blue fluorescent hybridization) is shown in g-h’’’. Details of the squared areas in a, c, e and g are shown in b-b’’, d-d’’, f-f’’, and h-h’’’. GFP cells of BSTM do not express GAD67 (b-b’’), but do express VGLUT2 (insert in d-d’’ and f-f’’); thus, they are glutamatergic. Triple labeling shows that the vast majority of the GFP cells in BST also express Foxg1 and VGLUT2 (insert in h-h’’’). For abbreviations see list. Scales: a, c, e, g = 200 µm; b, d, f = 50 µm (applies to b-b’’, d-d″, f-f’’); h = 50 µm (applies to h-h’’’).

**Figure 12.**
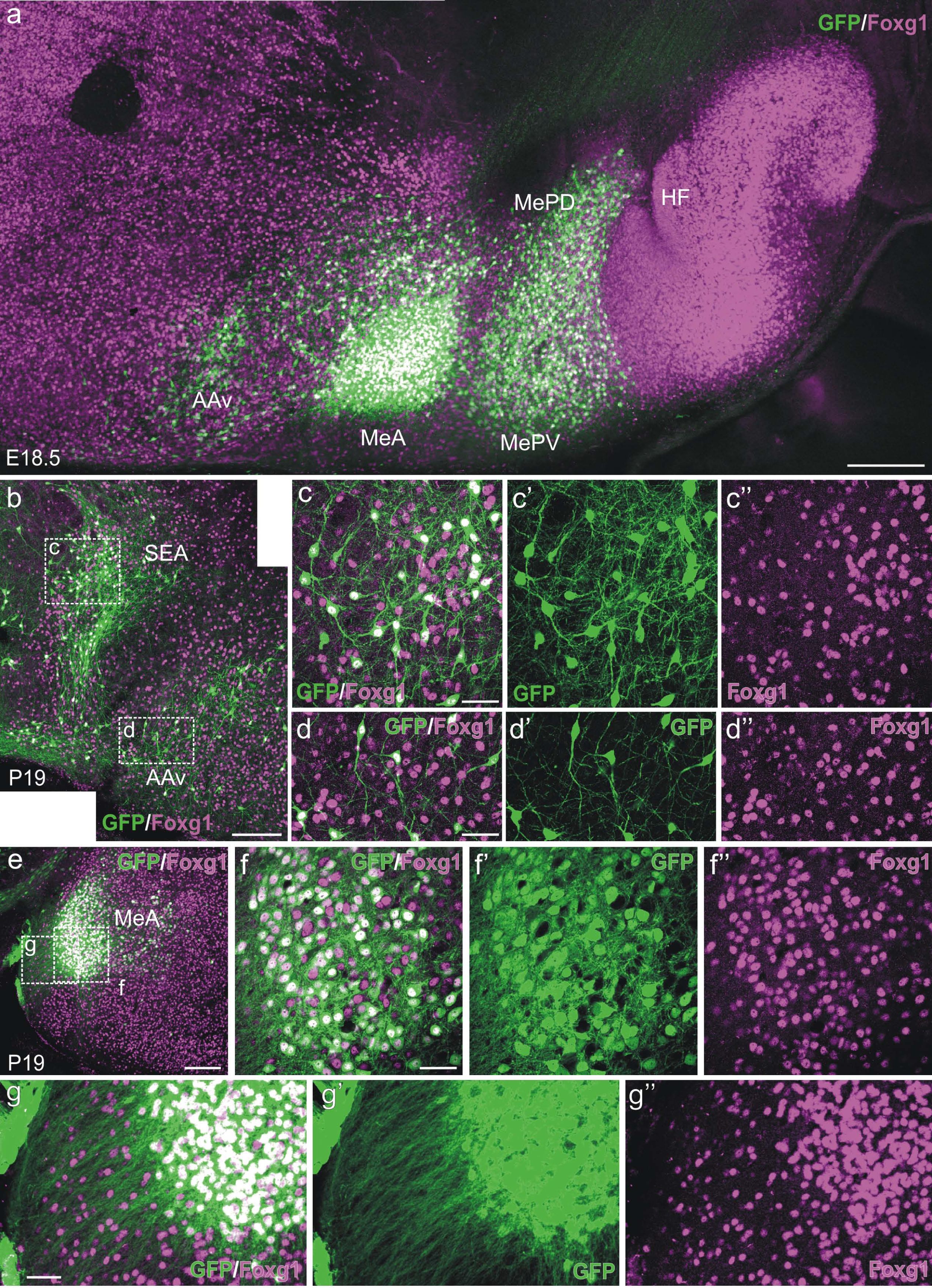
Confocal images of sagittal (a) and frontal (b-g’’) sections of E18.5 and P19 Otp-eGFP mice, at the level of the medial amygdala, anterior amygdala and sublenticular extended amygdala, double labeled for GFP (green) and Foxg1 (magenta) (a and b show photomontages). Details of the areas squared in b and e are shown in c-c’’, d-d’’, f-f’’ and g-g’’. Note the high density of GFP cells in MeA (striking) and in MeP (a bit less dense). The vast majority of these coexpress Foxg1. Double-labeled cells are also densely packed in a subdivision of the sublenticular extended amygdala (b and detail in c-c’’). In contrast, double labeled cells are more dispersed in the anterior amygdala (AAv, a, d-d’’). Note the labeled axons exiting the MeA in g-g’. For abbreviations see list. Scales: a, b, e = 200 µm; c, d, f, g = 50 µm (applies to c-c’’, d-d’’, f-f’’, g-g’’).

**Figure 13.**
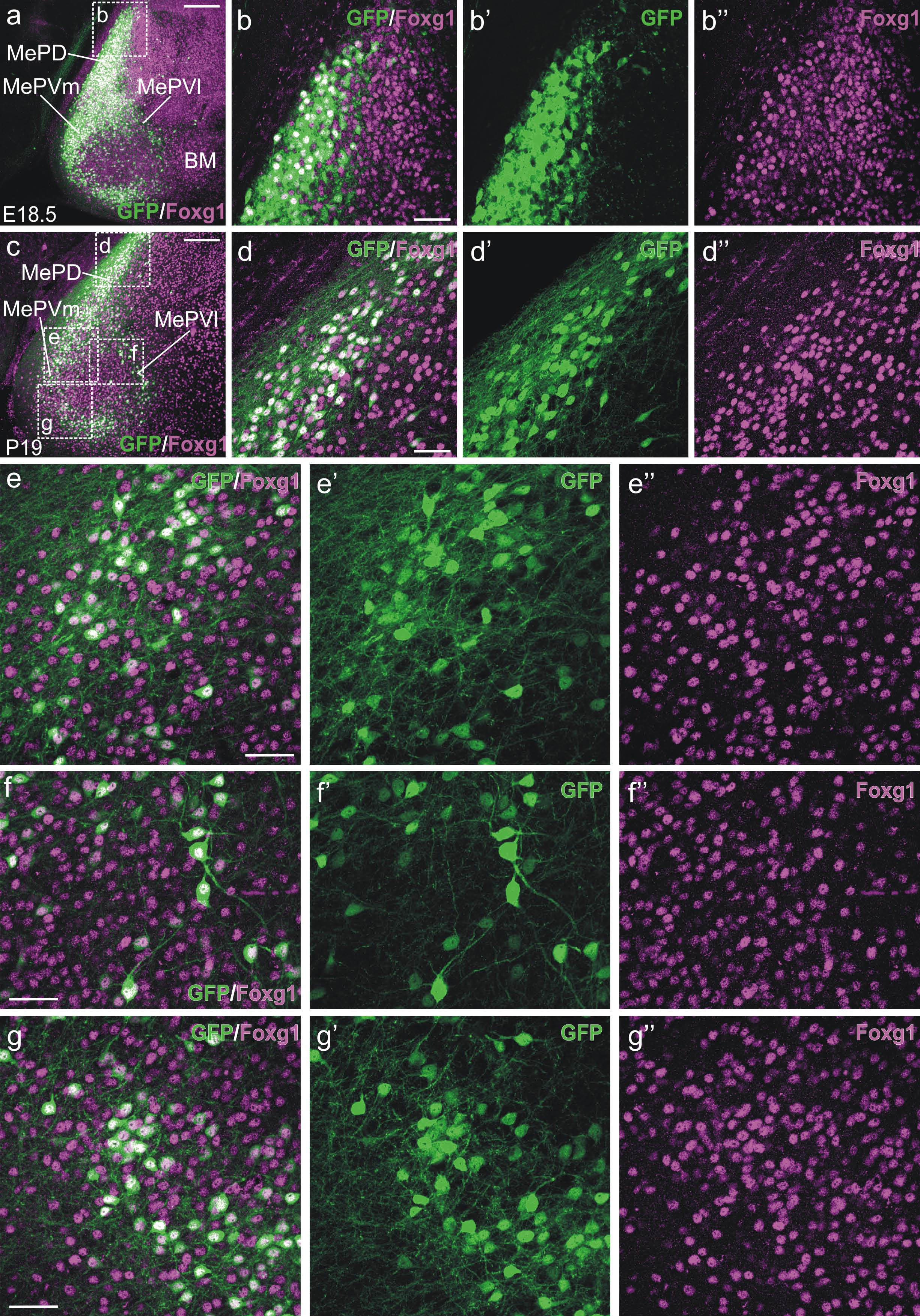
Confocal images of frontal sections of E18.5 and P19 Otp-eGFP mice, at the level of the posterior medial amygdala, double labeled for GFP (green) and Foxg1 (magenta). Details of the areas squared in a and c are shown in b-b’’, d-d’’, e-e’’, f-f’’ and g-g’’. Note the high density of GFP cells in the medial part of MePD (a-d’’). The vast majority of these co-express Foxg1 (d-d’’). In contrast, the lateral part of MePD is rich in Foxg1-only cells. In MePV, GFP cells (most co-expressing Foxg1) tend to concentrate in a ring-like area, surrounding a central area with dispersed GFP cells (a, c). Nevertheless, the density of double labeled cells is higher in the medial branch of the ring (e-e’’). For abbreviations see list. Scales: a, c = 200 µm; b, d, e, f, g = 50 µm (applies to b-b’’, d-d’’, e-e’’, f-f’’, g-g’’).

#### Preoptic region

As noted above, the TOH produced a part of the preoptic region, while another part derives from the telencephalic subpallium. Our analysis of the Otp-lineage cells (GFP-labeled) in combination with Foxg1 in this region throughout development helped us to distinguish between the derivatives of both divisions. At E18.5, double-labeled cells accumulated ventrally, in the anteroventral preoptic nucleus (POAV), extending from here into a ventral part of the lateral preoptic area, which also contained large amounts of double-labeled cells (**Figs. 9a-d; 10a**,**c**). These findings were also confirmed at P19 (**Fig. 9**). We interpret that this ventral preoptic region rich in GFP/Foxg1 double-labeled cells is derived from the terminal TOH. Nevertheless, this TOH part of the preoptic region also contained Fogx1-only and some scattered GFP-only cells, mostly intermingled between double-labeled cells (**Fig. 9**). Moreover, there was a group of GFP-only cells at the marginal zone of the lateroventral preoptic region (**Figs. 9e; 10a**,**b**), which appeared to correspond to a dorsal extension of the terminal prosomeric division of the supraoptic hypothalamic nucleus (SOt), as seen in sagittal sections; **Figs. 8b; 10a**,**b**). Another interesting observation is a massive GFP-labeled fiber tract apparently projecting from the TOH region of the preoptic region to the septum (**Fig. 8c, arrow**). The exact origin of these labeled fibers was unclear, and likely includes different TOH-derived cell subgroups as well as GFP cells of the hypothalamus. Fibers in the septum gave rise to a remarkable innervation in the lateral part, which was particularly dense in the lateroventral subnucleus. Dorsally, at precallosal levels, labeled fibers of the septum were in continuity with a thin cingular fascicle observed in the medial aspect of the cortex.

In contrast, at E18.5 and P19, dorsal parts of the preoptic region primarily contained Foxg1-only labeled cells, and represents the previously described subpallial preoptic subdivision, expressing Nkx2.1 and Shh in the vz during embryonic development (i.e., this is the subpallial, telencephalic part of the preoptic region) (**Fig. 9**). This region also included a subpopulation of GFP/Foxg1 double-labeled cells, some of which formed remarkable groups of cells along the periventricular preoptic nucleus and in the dorsolateral preoptic area (**Fig. 9**). This subpallial part of the preoptic region rich in Foxg1-only cells, in addition to having a subset of GFP/Foxg1 double-labeled cells (possibly with TOH origin), also contained a few GFP-only labeled cells (**Fig. 9**).

#### Extended amygdala

The extended amygdala was rich in Foxg1-only expressing cells (as typical in telencephalic structures), but part of its medial subdivision (including parts of medial BSTM and medial amygdala) was primarily populated by densely-packed GFP cells, the vast majority of which co-expressed Foxg1 (**Figs. 10-14**). These subdivisions of the medial extended amygdala rich in double-labeled cells likely derive from the peduncular TOH, and were distinguished by a dense GFP-labeled neuropil. In contrast, the central extended amygdala (central amygdala and lateral BST) did not contain GFP-labeled cells, but showed a weak or moderate labeled neuropil.

##### BSTM and sublenticular extended amygdala

Careful analysis of frontal and sagittal sections at E18.5 and P19 showed the following gradients from anterior to posterior, and from medial to lateral parts of BSTM. The anterior part of BSTM was mostly devoid of GFP-labeled cells (with a few exceptions possibly related to tangential migrations) (**Fig. 10a**,**a’**). Double cells were primarily located in the posterior part of the BSTM (**Figs. 10a**,**b-h’’; 11h-h’’**). Many of these cells were condensed in a narrow medial stripe of the principal subnucleus (or posteromedial BSTM), while they showed a dispersed organization more laterally. The posterior BSTM also contain many Foxg1-only labeled cells, with the exception of the medial stripe rich in double-labeled cells (**Fig. 10e-f’’**). At more caudal levels, the distribution of double-labeled cells in the posterior BSTM was similar, but an additional group of GFP-only labeled cells could be identified medially to the posteromedial BSTM (**Fig. 10g**). Even more caudally, both the GFP/Foxg1 double-labeled and the GFP-only cells of the posterior BSTM became continuous with similar double-labeled and single-labeled cells of the supracapsular BST. While the double-labeled cells clearly derived from peduncular TOH, our observations suggested that the GFP-only labeled cells of both posterior BSTM and supracapsular BST might derive from the peduncular SPV core (as the latter gave rise to a dorsal spike caudally adjacent to the TOH part of BSTM, corresponding to the previously named hypothalamic part of BSTM) (**Fig. 1d**).

The GFP labeled cells of the posterior BSTM were also continuous laterally with those found along the sublenticular extended amygdala, and the latter were in continuity with a large group found in the medial amygdala, all forming part of the TOH radial domain (**Figs. 7, 12**). Analysis at E18.5 and P19 showed that the vast majority of these cell populations were Foxg1/GFP double-labeled (**Fig. 12**).

##### Medial amygdala

Both the anterior (MeA) and posterior (MeP) subnuclei of the medial amygdala contained abundant GFP/Foxg1 double-labeled cells (**Figs. 12, 13**). In the MeA, double-labeled cells formed a dense group in the anterodorsal division (**Fig. 12**). In the MeP, the double-labeled cells occupied the medialmost aspect of the posterodorsal nucleus (MePD), while the lateral two-thirds were primarily occupied by Foxg1-only labeled cells (**Figs. 13**). In the posteroventral nucleus (MePV), double-labeled cells primarily organized forming a ring around a center rich in Foxg1-only cells with scattered double-labeled cells (**Fig. 13a**,**c**). The medial part of the ring was particularly rich in double-labeled cells (MePVm in **Fig. 13a**,**c**).

##### Other parts of the amygdala

Subsets of double-labeled cells appeared to split off from the main radial group, giving rise to tangentially dispersed cell subpopulations found in other parts of the medial extended amygdala, such as the anteroventral amygdala (AAv, in the subpallial amygdala) and the intraamygdaloid BST (BSTia) (**Figs. 7c; 8a; 12a**,**b**). In addition, some cells were also found in the pallial amygdala, including the anterior cortical amygdalar area and the basomedial nucleus of the basal amygdalar complex (**Fig. 7c**).

##### Phenotype

To know the phenotype of the Foxg1/GFP double-labeled cells in the medial extended amygdala, we carried out double labeling for GFP and either VGLUT2 or GAD67, and triple labeling for GFP, Foxg1 and either VGLUT2 or GAD67 at P19 (**Figs. 11, 14**). First, our double-labeling data showed that GFP cells in BSTM and medial amygdala did not express GAD67 (**Figs. 11a**,**b-b’’; 14a**,**b-b’’**), but instead expressed VGLUT2 (**Figs. 11d-d’’**,**f-f’’; 14f-f’’**,**h-h’’**,**i-I’’**). Moreover, triple labeling showed that the vast majority (if not all) of Foxg1/GFP double-labeled cells in the medial amygdala and BSTM expressed VGLUT2 and were, therefore, glutamatergic (**Figs. 11h-h’’’; 14g-g’’’**). In contrast, while many Foxg1-only cells in these regions were GABAergic (i.e., co-expressed GAD67), none of the GFP/Foxg1 double-labeled cells contained GAD67.

**Figure 14.**
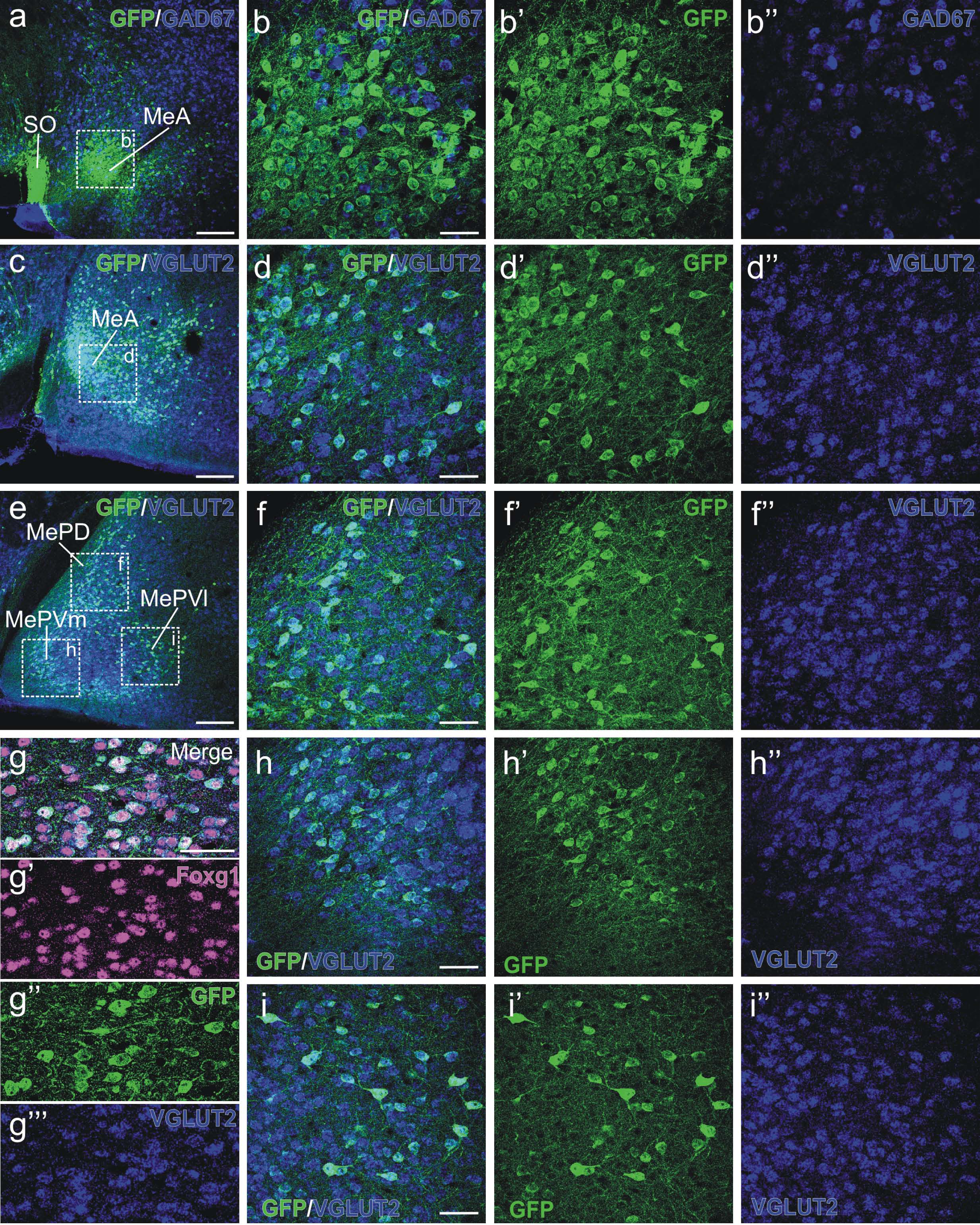
Confocal images of details of frontal brain sections of P19 Otp-eGFP mice, at the level of the BSTM, double labeled for GFP (green immunofluorescence) and either for GAD67 or VGLUT2 (blue fluorescent in situ hybridization; a-b’’ GAD67; c-f’’ and h-I’’ VGLUT2). Triple labeling for GFP (green), Foxg1 (magenta) and VGLUT2 (blue fluorescent hybridization) is shown in g-g’’’. Details of the areas squared in a, c and e are shown in b-b’’, d-d’’, f-f’’, h-h’’ and i-i’’. GFP neurons of the medial amygdala do not express GAD67 (b-b’’), but do express VGLUT2 (d-d’’, f-f’’, h-h’’, i-i’’). Triple labeling shows that the vast majority of these GFP glutamatergic cells also express Foxg1, as shown in a detail in the medial part of MePV (g-g’’’). For abbreviations see list. Scales: a, c, e = 200 µm; b, d, f, g, h, i = 50 µm (applies to b-b’’, d-d’’, f-f’’, g-g’’’, h-h’’, i-i’’).

##### Neuropil and projections

In general, the areas of the medial extended amygdala containing GFP labeled cells also show abundant labeled neuropil (**Figs. 7, 8, 10h; 12, 13**). We studied this in detail and distinguished some axonal projection thanks to the Golgi-like labeling of cell bodies, dendrites and axons in the Otp-eGFP transgenic mice. In the medial amygdala, labeled axonal terminals were often observed on labeled cell bodies and proximal dendrites. In addition, we also observed some terminals on non-labeled cells (**Fig. 13e’**,**g’**).

Notably, GFP-labeled cells of the medial amygdala gave rise to two projection fiber tracts (**Figs. 8, 12**):

(1) The dorsal amygdalofugal tract or stria terminalis (**Fig. 8a**), projecting to the BST. This might reach other known targets of the medial amygdala in the rostromedial telencephalon, such as nucleus accumbens, the Calleja islands, and the septum, all of which showed moderate to dense innervation by GFP-labeled fibers (**Fig. 7c; 8c**).

(2) The ventral amygdalofugal tract (**Fig. 12g-g’; 13d**), descending through the ansa lenticularis. These labeled axons might reach several known targets of the medial amygdala in the hypothalamus, paraventricular thalamus, and periaqueductal gray, all of which were innervated by GFP-labeled fibers (**Fig. 15-17**). Since the vast majority of the GFP cells of the medial amygdala coexpress Foxg1, at least part of the GFP-labeled innervation observed in the hypothalamic, thalamic and brainstem targets likely arise in double-labeled cells of the amygdala. Nevertheless, at least part of the GFP-labeled innervation of these areas may originate from GFP-only cells of the hypothalamus, which also project to some of the same targets.

**Figure 15.**
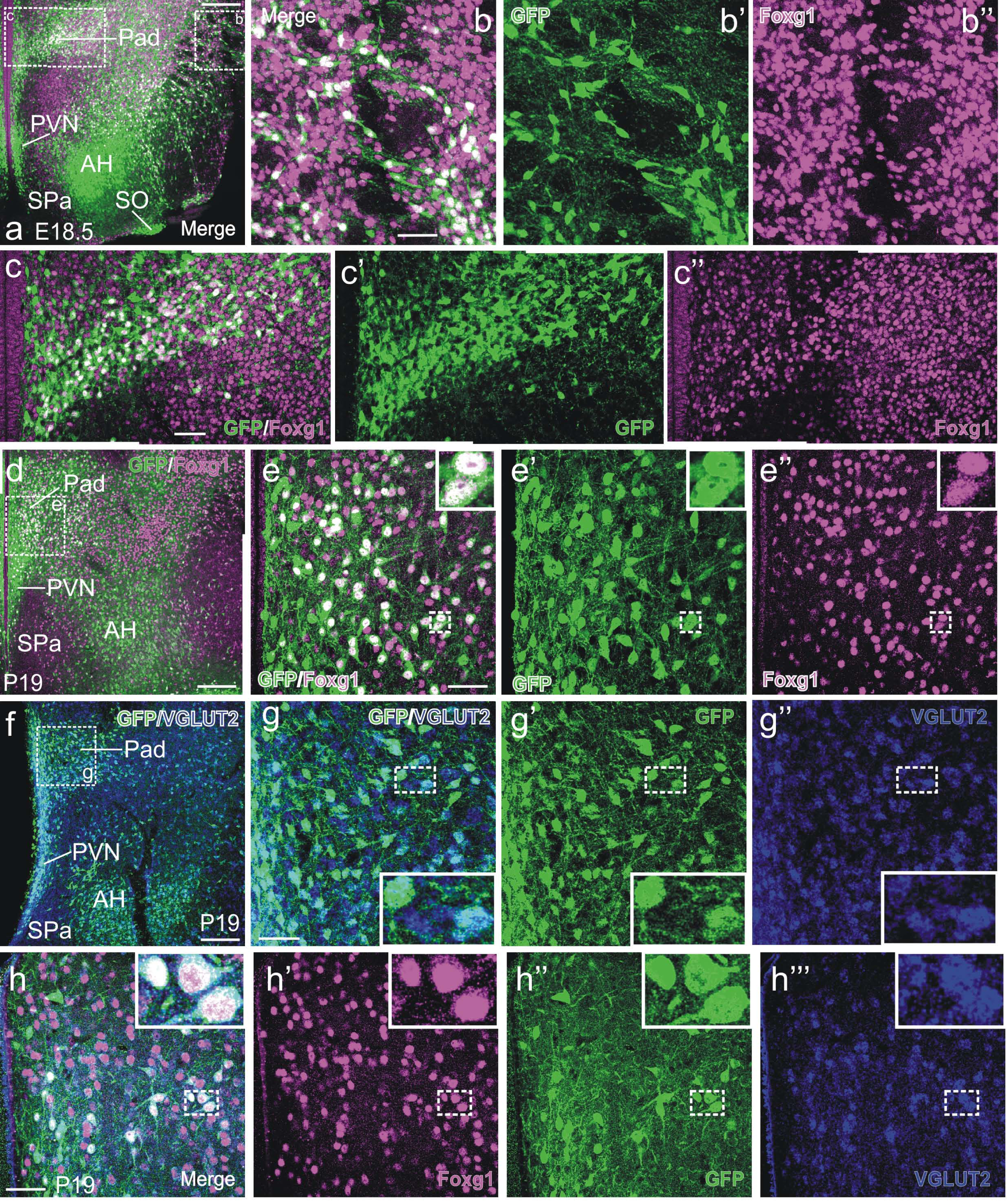
**a-e’’:** Confocal images of frontal sections of E18.5 and P19 Otp-eGFP mice, at the level of the dorsal paraventricular nucleus (Pad), double labeled for GFP (green) and Foxg1 (magenta). Details of the areas squared in a and d are shown in b-b’’, c-c’’ and e-e’’ (c-c’’ show photomontages). The Pad, part of TOH, contains a high density of GFP cells that are double labeled for Foxg1 (inserts in e-e’’). In contrast, the main part of the paraventricular nucleus (PVN) is rich in GFP single labeled cells. **f-g’’:** Confocal images of a frontal brain section of P19 Otp-eGFP mice, at the level of Pad, double labeled for GFP (green immunofluorescence) and VGLUT2 (blue fluorescent in situ hybridization). A detail of the area squared in f is shown in g-g’’. These images show that the GFP cells of Pad express VGLUT2 (insert in g-g’’). **h-h’’’:** Detail of a brain frontal section of a P19 Otp-eGFP mouse, showing triple labeling for GFP (green), Foxg1 (magenta) and VGLUT2 (blue fluorescent hybridization) in cells of Pad (inserts). For abbreviations see list. Scales: a, d, f = 200 µm. b, c, e, g, h = 50 µm (applies to b-b’’, c-c’’, e-e’’, g-g’’, h-h’’’).

#### Paraventricular-supraoptic complex

According to our data, the peduncular TOH seems to produce cell groups previously attributed to the hypothalamus, such as a dorsal part of the so-called paraventricular nucleus, as well as dorsal parts of the so-called anterior and lateral hypothalamic areas, all of which were rich in GFP/Foxg1 double-labeled cells (**Fig. 15**). In the latter areas, we observed remarkable radially-oriented rows of double-labeled cells that extended near the surface. These cells near the surface had a tangential disposition, and were in continuity with double-labeled cells of the episupraoptic nucleus (ESO, **Fig. 16b**,**c-c’’**). This suggested that double-labeled cells found in the ESO (a nucleus located more ventrally, in SPV core) might originate in TOH.

**Figure 16.**
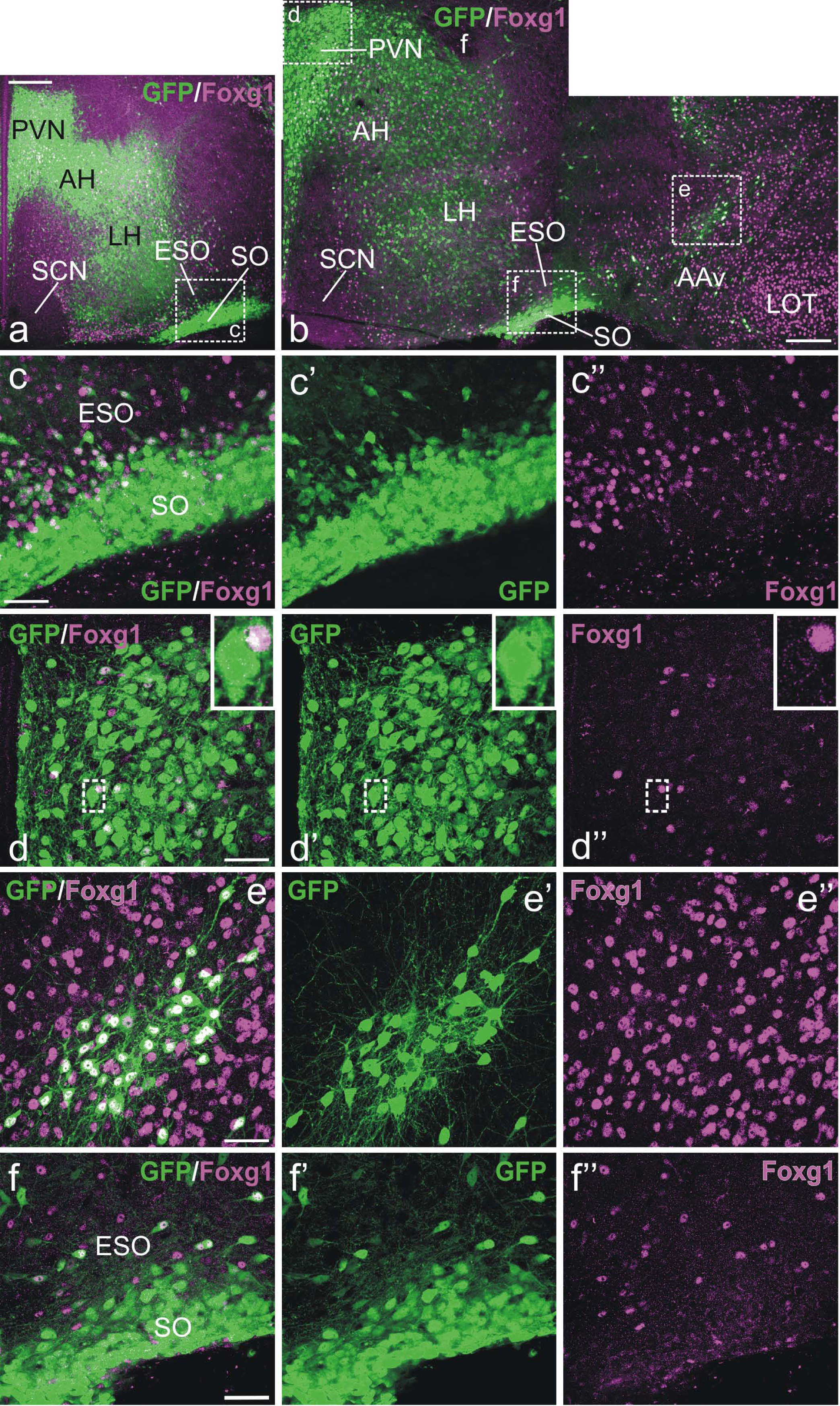
Confocal images of frontal sections of E18.5 (**a**) and P19 (**b-f’’**) Otp-eGFP mice, at the level of the principal parts of the paraventricular (PVN) and supraoptic (SO) nuclei, double labeled for GFP (green) and Foxg1 (magenta). b is a photomontage. Details of the areas squared in a and b are shown in c-c’’, d-d’’, e-e’’, f-f’’. Both PVN and SO contain a high density of GFP single labeled neurons. Nevertheless, these nuclei, part of SPV core, also contain a few double labeled cells (insert in d-d’’) as well as a few Foxg1 single labeled cells. Curiously, while GFP-only cells of PVN have a large soma (magnocellular), those double-labeled (GFP/Foxg1) are parvocellular. See text for more details. For abbreviations see list. Scales: a, b = 200 µm. c, d, e, f = 50 µm (applies to c-c’’, d-d’’, e-e’’, f-f’’).

In contrast to the dorsal region, the rest of the paraventricular and supraoptic nuclear complexes were rich in GFP-only cells and appeared to derive from SPV core. In the peduncular part of this region, GFP-only labeled cells were located periventricularly, in the central/ventral parts of the paraventricular hypothalamic nucleus (**Fig. 16**). More laterally, it included central parts of the anterior and lateral hypothalamic areas, and the main supraoptic nucleus was found at the marginal zone (**Fig. 16a**,**b**). All of these nuclei and areas were rich in GFP-only cells also included minor subsets of Foxg1-only cells and GFP/Foxg1 double-labeled cells (**Fig. 16c**,**d**). The ESO, located deep to the main supraoptic nucleus, contained subpopulations of the three types of neurons at E18.5 and P19 (**Fig. 16c**). We noticed the massive GFP-labeled projections from the supraoptic and paraventicular complexes to the median eminence (**Figs. 8c, 17c**). These labeled fibers seem to primarily originate in the SPV core (hypothalamic) subdivision of the complex, but we cannot discard an input from GFP cells of TOH.

Regarding the terminal prosomeric part of SPV core, this was very narrow at periventricular levels (as seen in sagittal sections), and it contained the terminal part of the supraoptic nucleus at its subpial surface (marginal zone) (**Figs. 1, 10**). Cells of this group were GFP-only labeled, and sagittal sections showed a subpial continuum of these cells from the main group in terminal SPV core into more dorsal divisions, reaching TOH and subpallial preoptic levels (**Fig. 10a**,**c**).

#### Subparaventricular and basal hypothalamic regions

As noted above, our data showed the presence of Otp-lineage cells (GFP labeled) in subparaventricular and basal parts of the hypothalamus, and some of them were GFP/Foxg1 double-labeled cells, suggesting a TOH origin (**Figs. 1, 5, 8, 17**). The subparaventricular region of the alar hypothalamus included a group of tangentially oriented double-labeled cells near the surface at peduncular prosomeric levels (not shown). In contrast, at terminal levels, this region was almost devoid of double-labeled cells. This was very evident in the medial aspect, where the suprachiasmatic nucleus is located (**Figs. 1c; 8c**). Lateral to it, we observed mainly non-overlapping subpopulations of Foxg1-only or GFP-only labeled cells.

**Figure 17.**
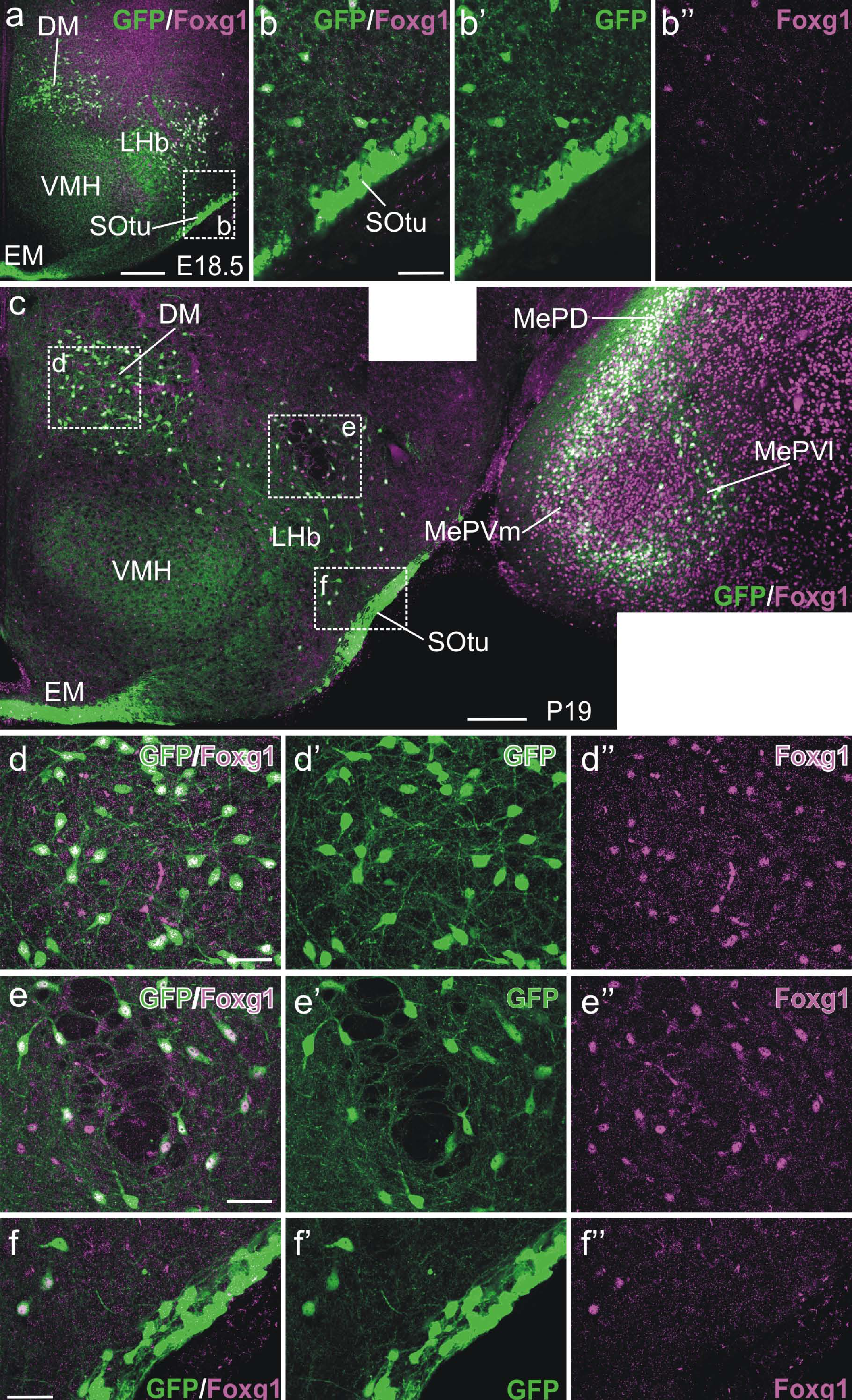
Confocal images of frontal sections of E18.5 (**a-b’’**) and P19 (**c-f’’**) Otp-eGFP mice, at the level of the basal hypothalamus, double labeled for GFP (green) and Foxg1 (magenta). c is a photomontage. Details of the areas squared in a and c are shown in b, d-d’’, e-e’’ and f-f’’. A subpopulation of double labeled cells is seen in the retrotuberal region, intermingled with GFP or Foxg1 single labeled cells (d-d’’ and e-e’’). In contrast, the tuberal supraoptic nucleus (SOtu) is rich in GFP single labeled cells (b-b’’, f-f’’). See text for more details. For abbreviations see list. Scales: a, c = 200 µm. b, d, e, f = 50 µm (applies to b-b’’, d-d’’, e-e’’, f-f’’).

The basal hypothalamus contained important subpopulations of GFP labeled cells from early stages of development (**Fig. 5b**). A large group of these cells was located in the peduncular basal hypothalamus, in a superficial area of the retrotuberal region devoid of Nkx2.1; moreover, the majority of these GFP cells (but not all) did not co-express Nkx2.1, suggested a non-basal hypothalamic origin (**Fig. 5b**). Throughout development, this superficial group of the retrotuberal hypothalamus was in clear continuity with GFP cells of the alar hypothalamus (**Fig. 1**). From late embryonic stages, GFP cells of the retrotuberal group were organized in medial and lateral subgroups, and both contained subpopulations of Foxg1/GFP double-labeled cells (**Fig. 17a**,**c**). This region also contained subsets of Foxg1-only cells, intermingled with the double-labeled cells. In addition, a distinct group of GFP-only cells was observed at the subpial surface of the retrotuberal region, which extended into the marginal zone of the tuberal region (in the terminal prosomeric division). This corresponded to the tuberal supraoptic nucleus (**Fig. 17a-c**). The fact that cells of this group were GFP-only labeled suggested an origin in SPV core.

## Discussion

### The TOH in the context of the forebrain building plan

The vertebrate brain develops and is organized according to a common building plan (Bauplan or morphoplan), which means that during development it gradually subdivides into a shared series of longitudinal (i.e. dorsoventral) and transversal compartments in response to external and internal morphogenetic signals (Nieuwenhuys and Puelles, 2016). Each major compartment is additionally subdivided in smaller domains, so that at the end the neural tube consists of a number of subdivisions comparable across species, each one occupying a specific topological position in the morphospace (Nieuwenhuys, 1998; Puelles and Medina, 2002).

Importantly, the genes that govern the specification and other early steps of the formation of each subdivision are highly conserved in evolution (Puelles et al., 2000; Moreno and González, 2006; Medina and Abellán, 2009; Abellán and Medina, 2009; Abellán et al., 2014; Desfilis et al., 2018). Thus, at early developmental stages, homologous embryonic domains share an identical topological position within the Bauplan and a highly similar molecular signature (Puelles and Medina, 2002; Puelles and Ferran, 2012; Nieuwenhuys and Puelles, 2016). An important consequence of this is that neurons derived from each compartment are characterized by combinatorial expression of highly conserved developmental regulatory genes, including genes encoding transcription factors and other proteins involved in regulation of early developmental processes; this early molecular signature is going to determine later aspects like cell migration, axonal pathfinding, and maturation (Butler and Tear, 2007; Chedotal and Richards, 2010; Williams et al., 2010; discussed by Medina et al., 2017a,b). For this reason, knowledge on the major divisions and subdivisions that conform the Bauplan is crucial to understand the diversity of neurons and the connectivity patterns found in the mature brain. In fact, this information can be decisive to understand the logic behind the huge amount of connections found in the mature brain (García-López et al., 2008; Medina et al., 2011, 2017a,b; Sokolowski and Corbin, 2012; Hassan and Hiesinger, 2015). In this context, the finding of a new molecular domain at the transition between telencephalon and hypothalamus is highly relevant, because it opens new venues to study the distribution, chemical phenotype and connections of the neurons produced there. A similar domain, with identical topological position and molecular signature (expression of both Foxg1 and Otp), has recently been found in the zebrafish (Affaticati et al., 2015), suggesting that it may represent a common feature of the Bauplan of the vertebrate brain, a question that needs further evaluation by studying additional species from different groups of vertebrates. The new division found so far in mouse (present results) and zebrafish (Affaticati et al., 2015) is located within the alar region of the secondary prosencephalon, and -as seen in sagittal sections-extends through peduncular and prepeduncular (terminal) segmental subdivisions (as defined in the latest version of the prosomeric model by Puelles and Rubenstein, 2015; Ferran et al., 2015; see also scheme in **Figure 1**). The peduncular prosomeric subdivision can also be distinguished by its Rgs4 expression (Ferran et al., 2015), but according to our data, Rgs4 is highly expressed only in the peduncular part of the hypothalamus (in SPV core; see results), but not in TOH. This new division appears to produce a region previously considered part of the preoptic area and adjacent hypothalamus, as well as part of the optic vesicle (Affaticati et al., 2015; for this reason, the latter authors called the new domain the ‘optic recess region’). The mouse TOH and the comparable division of zebrafish (Affaticati et al., 2015) show overlapping expression of both GFP/Otp and Sim1. However, our results in mouse show that coexpression only occurs in some postmitotic cell subpopulations. In addition, our holistic analysis in the mouse, using several sectioning planes combined with comparisons to Foxg1 and radial glia, demonstrated that the TOH is a truly radial histogenetic division, which gives rise to a part of the medial extended amygdala rich in GFP/Foxg1 double-labeled cells. Therefore, our study leads to a reinterpretation of the telencephalon and the hypothalamus, as discussed in next section. In addition, our study suggests that the TOH gives rise to several GFP/Foxg1 double-labeled cell subpopulations that migrate tangentially to some parts of the telencephalon and the hypothalamus. Like in zebrafish, this domain in mouse seems to relate to an anterior part of the eye vesicle and pedicle, which shows Foxg1 expression and is laterally adjacent to the rest of TOH (Dou et al., 1999; Crossley et al., 2001).

### Distinction of TOH and its derivatives from those of adjacent areas

We took advantage of the transgenic Otp-eGFP mouse with permanent labeling of Otp-lineage cells in order to investigate the phenotype of TOH-derived cells, which could help us to distinguish them from those derived from adjacent divisions. Our results demonstrate that TOH-derived cells truly co-express GFP and Foxg1 and seem to originate from Foxg1 progenitors lining the interventricular foramen (Monro) and the immediately adjacent third ventricle. Cells derived from this domain can thus be distinguished from those derived from progenitors in the telencephalon (expressing only Foxg1 but not GFP) as well as from progenitors in the immediately adjacent paraventricular hypothalamic region (in particular, the SPV core, producing cells that express GFP but not Foxg1). This feature helped us to distinguish TOH-derived cells from those of adjacent areas, and study their dispersion throughout development.

Based on expression of Otp alone (i.e., compared to other transcription factors such as Dlx5, but without considering Foxg1), the TOH was previously included as part of a subdivision of the optoeminential region, initially called the supraopto-paraventricular hypothalamic (SPV) domain (Puelles and Rubenstein, 2003), and later simply called paraventricular hypothalamic domain (Bardet et al., 2008; Morales-Delgado et al., 2011; Puelles et al., 2012; Puelles and Rubenstein, 2015; Ferran et al., 2015; see also Díaz et al., 2015). This Otp-expressing domain was described to co-express Sim1 but not Dlx2/5, and to be bounded by Dlx2/5-expressing domains of the subpallium (dorsally), subparaventricular hypothalamic region (ventrally) and prethalamus (caudally) (Simeone et al., 1994; Acampora et al., 2000; Puelles and Rubenstein, 2003; Bardet et al., 2008; Puelles et al., 2012; Morales-Delgado et al., 2011). An Otp-expressing domain occupying an identical topological position has also been found in birds (chicken: Bardet et al., 2008), reptiles (turtle: Moreno et al., 2012; long-tailed lizard: Abellán et al., 2013), amphibians (frog: Domínguez et al., 2013) and actinopterygian fishes (zebrafish: Biechl et al., 2017). In agreement with previous studies in turtles (Moreno et al., 2012) and frogs (Domínguez et al., 2013), the Otp-expressing SPV domain in mouse can also be distinguished from the telencephalic subpallium, the subparaventricular hypothalamic domain and the prethalamus by the expression patterns of Nkx2.1 and Islet1: the SPV appears as a negative gap regarding the expression of these transcription factors (present results). According to the prosomeric model, the dorsalmost part of SPV directly abuts the telencephalic subpallium, except at its caudal end, where it shows a spike-like extension into the pallial amygdala (Puelles and Rubenstein, 2003; Puelles et al., 2012). This SPV-related spike extending into the amygdala is present not only in mouse, but also in species of different vertebrates (Bardet et al., 2008; Moreno et al., 2012; Biechl et al., 2017). However, our data show that the classical SPV is subdivided into the TOH (co-expressing GFP and Foxg1) and the SPV core (expressing GFP, but not Foxg1; also expressing Rgs4 in the peduncular division). Our proposal of TOH as a partition of classical SPV requires a reinterpretation of the relation between hypothalamus and telencephalic subpallium, since this is not direct, but through the TOH. The latter division, but not the hypothalamus, is the one directly abutting the subpallium. Regarding the spike-like extension of the Otp expression domain, our results show that its main part belongs to TOH (and is GFP/Foxg1 double-labeled), except for a narrow GFP-only band on the caudalmost side of the spike that relates to part of the supracapsular BST.

As shown by migration assays in mouse, the classical SPV was assumed to be a hypothalamic source of cells tangentially migrating to the medial extended amygdala (Soma et al., 2009; García-Moreno et al., 2010; Bupesh et al., 2011a; discussed by Medina et al., 2011, 2017a). The transcription factor Otp, expressed in postmitotic cells, was found to be essential for cell migration across the hypothalamo-telencephalic boundary (García-Moreno et al., 2010) and for differentiation of neuroendocrine cell lineages (Acampora et al., 1999, 2000; Wang and Lufkin, 2000). Based on our double-labeling studies in Otp-eGFP animals, it is likely that the vast majority of the cells migrating to the medial extended amygdala (medial amygdala and BSTM) and preoptic area in those experiments originate in the TOH, since most of them contain both Foxg1 and GFP. Moreover, our holistic analysis of the TOH radial domain involves a reinterpretation of the cell migrations to the medial extended amygdala, since a large part of them appear to occur radially within the TOH. This includes the GFP/Foxg1 double-labeled cells that populate the posterior BSTM, part of the sublenticular extended amygdala, as well as the anterodorsal subnucleus and medial aspects of the posterodorsal and possibly posteroventral subnuclei of the medial amygdala. In contrast, only smaller subsets of TOH-derived double-labeled cells might migrate tangentially into subpallial parts of the preoptic region and extended amygdala (including the anterior BSTM, the ventral anterior amygdala or AAv, and lateral aspects of the posterior medial amygdala) as well as into pallial parts of the amygdala (including the basomedial nucleus and anterior cortical amygdalar area).

Previous studies have also found other sources of extratelencephalic cells for the telencephalon: the prethalamic eminence has been shown to produce groups of Lhx5- and Pax6-expressing neurons that enter the telencephalon and reach as far rostral as the accessory olfactory bulb, leaving different cell types along the way (Roy et al., 2014; Ruiz-Reig et al., 2017), including cells for the extended amygdala (Abellán et al., 2010; Bupesh et al., 2011b; Ruiz-Reig et al., 2017). However, Lhx5 cells of the extended amygdala can also be produced inside the telencephalon and in classical SPV (Abellán et al., 2010; Vicario et al., 2017), and in mouse Lhx5 appears to be expressed in Otp-lineage SPV-derived cells (García-Moreno et al., 2010). Thus, some areas of the telencephalon, including the extended amygdala, contain cell subpopulations from different extratelencephalic sources, in addition to those produced locally, within the telencephalon, but it is not always easy to distinguish between them. However, how do we define the telencephalon? It is becoming more and more evident that the classical definition of the telencephalon as a rostrodorsal forebrain region expressing Foxg1, divided into pallial and subpallial parts, is now insufficient. Our data on the TOH, together with some previously published results on the prethalamic eminence, show that these two domains form the caudal part of the classical telencephalon (Abellán et al., 2010; Ruiz-Reig et al., 2017; present results). Independent of this philosophical dissertation, our data allow distinction between the different cell subpopulations located in the ventral part of the classical telencephalon: cells born in the pallido-preoptic region of the subpallium express Foxg1 and different transcription factors such as Nkx2.1 and Islet1 (among others: Carney et al., 2010; present results); in contrast, cells derived from TOH are Foxg1/GFP (Otp-lineage) double-labeled, but the vast majority of them do not express Nkx2.1 or Islet1; cells coming from the SPV core (excluding TOH) express only GFP but not Foxg1; finally, cells that originate in the prethalamic eminence express neither GFP nor Foxg1. This allowed us to safely trace TOH-derived cells in the classical telencephalon, and distinguish them from those produced locally or having other origins.

TOH-derived cells do not only spread dorsally into the telencephalon, but also appear to spread ventrally into the hypothalamus. Subsets of double-labeled cells have been found in both the alar hypothalamus (including the paraventricular and subparaventricular domains) as well as in the basal hypothalamus (mainly in the retrotuberal region). If confirmed by migration assays, this would raise questions on the chemical phenotype and connections of these immigrant double-labeled cells (further discussion below), and on their functional relationships with the GFP-only cells in the paraventricular and supraoptic hypothalamic nuclei (among other).

### TOH as a new source of glutamatergic neurons for the medial extended amygdala

Our results agree with those of García-Moreno et al. (2010) on the presence of a large subpopulation of Otp-lineage cells in the BSTM and medial amygdala, and on the glutamatergic phenotype of these cells (based on expression of VGLUT2). None of these cells express GAD67, a marker of GABAergic cells. However, our data show that the vast majority of these cells co-express Foxg1 and originate in the TOH. Our results also provide interesting information on the specific location of the TOH-derived (GFP/Foxg1-double-labeled) glutamatergic cells of the BSTM and medial amygdala, which allows comparisons with the Foxg1-only cell populations (most of which are GABAergic) (present results), as well as with previous studies on the connections and/or functions of the glutamatergic cells in these nuclei.

Regarding the anterior division of the medial amygdala, densely-packed GFP/Foxg1 cells are primarily located in the anterodorsal subnucleus and are intermingled with a few Foxg1-only cells. With respect to the posterior division, double-labeled cells mainly concentrate medially in the posterodorsal subnucleus (MePD), while in the posteroventral subnucleus (MePV) they primarily organize forming an external cellular ring surrounding an internal core rich in Foxg1-only cells; the inner core only contains a few dispersed double-labeled cells. However, also in MePV, the majority of the double-labeled cells concentrate in the medial aspect of the ring (Fig. 13a,c). These subregions within MePD and MePV are also distinguished based on cytoarchitecture, as seen in Nissl staining (Canteras et al., 1995; see also **Fig. 13** in our material). In general, areas of MePD and MePV with high density of double-labeled cells contain few Foxg1-only cells. As noted above, the Foxg1/GFP double-labeled cells of the medial amygdala co-express VGLUT2 and are, therefore, glutamatergic. This abundant glutamatergic cell population with TOH origin is complementary to other glutamatergic cells of the medial amygdala that originate in the ventral pallium (Bupesh et al., 2011a; Abellán et al., 2013; Medina et al., 2017a), the ventrocaudal pallium (Medina et al., 2017a; Ruiz-Reig et al., 2018) and the prethalamic eminence (Abellán et al., 2013; Medina et al., 2017a; Ruiz-Reig et al., 2017). Regarding the Foxg1-only cells of the medial amygdala, most of them are GABAergic cells that likely originate in the subpallium (in the ventrocaudal pallidal/diagonal and commissural preoptic subdivisions; García-López et al., 2008; Carney et al., 2010; Bupesh et al., 2011a). Comparison of our data with previous results of our group (García-López et al., 2008) indicates that in the posterior medial amygdala TOH-derived cells tend to occupy a partially different position to those from pallidal/diagonal or preoptic origins: in MePD, TOH cells primarily concentrate medially, while pallido/diagonal cells (expressing Lhx6) spread more laterally, with only partial overlap (for Lhx6 see Fig. 12 in García-López et al., 2008); in MePV, while TOH cells primarily locate in a peripheric ring (but mostly medially), those derived from the commissural preoptic region (expressing Shh) also occupy the inner part (for Shh, see Fig. 10 in García-López et al., 2008). In addition, some Foxg1-only cells of the medial amygdala might be glutamatergic, likely representing the subpopulations of Lhx9 and/or Ebf3 cells that originate in the ventral/ventrocaudal pallium (Medina et al., 2017a; Ruiz-Reig et al., 2018); while ventral pallial cells tend to concentrate close to the surface, outside de ring of TOH-derived cells (García-López et al., 2008; Bupesh et al., 2011a; Medina et al., 2017a), those from the ventrocaudal pallium mainly occupy the lateral part of MePV ring (Ruig-Reig et al., 2018).

With respect to the BSTM, double-labeled cells primarily locate in its posterior division, and show a high concentration in the medialmost aspect of the principal subnucleus (also called posteromedial BSTM), but scattered cells are also found more laterally, including posterointermediate and posterolateral parts (see other details in the Results). In all of these subdivisions, GFP/Foxg1 double-labeled cells overlap with Foxg1-only cells. Our data show that while the GFP/Foxg1 double-labeled cells are glutamatergic (i.e. they express VGLUT2), Foxg1-only cells are GABAergic (they express GAD67). As in the medial amygdala, the Foxg1-only GABAergic cells of the posterior BSTM likely include subpallial cells derived from the pallidal/diagonal division (expressing Lhx6) and cells derived from the subpallial preoptic divisions (expressing Shh, Islet1, Dbx1 or Foxp2) (García-López et al., 2008; Bupesh et al., 2011a; Lischinsky et al., 2017; Vicario et al., 2017). In mouse, subpallial cells of the posterior BSTM seem to reach this nucleus by tangential migration, since this nucleus forms within the radial extension of TOH (present results). Based on this, it appears that only the anterior BSTM, but not posterior BSTM, would form within the radial histogenetic domain of the ventrocaudal pallidum (which partially modifies a previous proposal by Bupesh et al., 2011a).

What are the connections and functions of these different cell types of the medial extended amygdala? Here we will focus on the medial amygdala or medial nuclear complex, which connections have been studied in great detail (for example, Canteras et al., 1995; Dong et al., 2001). The projections course through two major tracts: (1) the dorsal amygdalofugal tract or stria terminalis, which include different subcircuits related to different cell subpopulations (Bupesh et al., 2011a; present results). Our data supports that this is a radially disposed tract connecting cells of the medial amygdala with those in the BSTM. Some axons of this tract split off to target other telencephalic structures like nucleus accumbens (mainly the shell) and the septum (Canteras et al., 1995). Part of the fibers also seem to separate to reach the medial preoptic region, while another group pass through the stria medullaris to reach the paraventricular thalamus (Canteras et al., 1995). (2) The ventral amygdalofugal tract; this tract courses through the ansa lenticularis (a classical name that refers to part of the internal capsule) to reach the hypothalamus (anterior, ventromedial and pre-mammillary hypothalamic areas) and some brainstem areas, including the central gray and the ventral tegmental area (Canteras et al., 1995). Each medial amygdala subnucleus shows a specific pattern of projections. However, since each major subnucleus of the medial amygdala includes a mixture of different cell subpopulations (reviewed by Medina et al., 2017a), it is not possible to distinguish the specific connections of distinct neuron subpopulations by way of classical tract-tracing experiments.

GABAergic neurons originate in the subpallium, but are heterogeneous since they include different subpopulations from distinct lineages with pallidal or preoptic origin, which can be distinguished by their distinct molecular signature (García-López et al., 2008; Hirata et al., 2009; Carney et al., 2010; Bupesh et al., 2011a; Abellán et al., 2013; Lischinsky et al., 2017). By way of genetic encoded axonal tracers, Choi et al. (2005) were able to study the connections of some neuron subpopulations of the medial amygdala, including the Lhx6-expressing neurons (with origin the ventrocaudal pallidal/diagonal domain; García-López et al., 2008; Bupesh et al., 2011a). Choi et al. (2005) showed that the Lhx6-expressing neurons of the medial amygdala are activated by exposure to sex-related odors and preferentially project to Lhx6-rich subdivisions of the BSTM and ventromedial hypothalamus, which are involved in sexual behavior (Choi et al., 2005). This suggests that this transcription factor may help to delineate a reproduction-related neural subcircuit during brain development, which is highly conserved in evolution (Medina et al., 2011; Abellán et al., 2013; 2017a; Sokolowski and Corbin, 2012). We previously suggested that a similar principle could be applied to other neuron subtypes of the medial amygdala, and that neurons with different origin might get engaged in different functional networks (Medina et al., 2011; Abellán et al., 2013; Medina et al., 2017a). This developmentally-based principle could offer light to explain preferential connections between neurons sharing a similar molecular signature located in a particular region or distributed across multiple regions of the brain, providing a foundation for the formation of distinct functional neural subcircuits or pathways (discussed by Medina et al., 2011; Abellán et al., 2013; Medina et al., 2017a,b). This could also explain recent single-cell transcriptomic results in the neocortex of adult animals, which show that distinct subsets of molecularly-identified pyramidal neurons, each with a particular spatial distribution, could underlie the connectivity patterns found in the mature brain (Tasic et al., 2018). While the latter study highlights the relevance of molecular taxonomy of brain cells in order to understand neuron connections and function (commented by Bhaduri and Nowakowski, 2018), here we make emphasis on developmental studies to explain the molecular diversity and location of different neurons, as well as to predict the connections of neurons with the same origin and molecular signature.

Although more data are needed on the specific connections of different amygdala neuronal subtypes, it now appears that subsets of GABAergic neurons of the medial extended amygdala with distinct molecular signature and origin are involved in different types of social interactions (Choi et al., 2005; Lischinsky et al., 2017). What about the glutamatergic cells of the medial extended amygdala? We discuss this in next section.

### A glutamatergic subcircuit of the medial extended amygdala and its role in social behavior

Like the GABAergic neurons, glutamatergic neurons of the medial extended amygdala also are heterogeneous (Choi et al., 2005; Abellán et al., 2013). Based on their embryonic origin, these neurons can originate at least in the ventral/ventrocaudal pallium, TOH, alar hypothalamus (SPV core) and prethalamic eminence (reviewed by Medina et al., 2011, 2017a; Abellán et al., 2013; Ruiz-Reig et al., 2017, 2018; present results). Similarly, glutamatergic cells of the BSTM seem to originate in TOH, hypothalamus and prethalamic eminence (reviewed by Medina et al., 2011, 2017a; Abellán et al., 2013; Vicario et al., 2017; present results). Nevertheless, our results in mouse based on triple labeling of GFP, Foxg1 and VGLUT2 showed that the majority of the glutamatergic cells of the medial amygdala and BSTM originate in TOH. What are the connections and function of TOH-derived glutamatergic cells?

Interestingly, a recent study by Hong et al. (2014) showed that pharmacogenetic activation of VGLUT2 positive neurons in MePD leads to marked reduction of social interactions and promote self-grooming, resembling some of the social deficits observed in mouse models of autism (Hong et al., 2014). The deficits were specific for social behavior, since interactions with novel objects were not altered in these animals. The VGLUT2 positive cells identified by Hong et al. (2014) show an identical position to the GFP/Foxg1/VGLUT2 identified in our study (particularly in MePD), suggesting they may originate in TOH.

In the study mentioned above, it was unclear how the asocial and self-grooming behaviors were achieved since the connections of the VGLUT2 positive neurons were unknown. Our results show that TOH-derived glutamatergic neurons of the medial amygdala display GFP-labeled axons that exit the nucleus by way of both the dorsal and ventral amygdalofugal tracts. Thus, they are projection neurons. This agrees with the finding of glutamatergic projections from the medial extended amygdala to the ventromedial hypothalamus (Choi et al., 2005; see also Kiss et al., 2011). Moreover, we observed abundant GFP-labeled neuropil (fibers and terminals) in many of the known targets of the medial amygdala in the BSTM, septum, ventral striatum (accumbens shell and Calleja islands), preoptic region, hypothalamus (alar and basal parts), paraventricular thalamus, periaqueductal gray and ventral tegmental area (for the connections see Canteras et al., 1995). Some of the innervated areas also include GFP/Foxg1 double-labeled cells (such as the preoptic area and the hypothalamus), which might also derive from TOH. Based on the developmental-based principle for establishment of connections explained above, we predict the existence of preferential connections between the GFP/Foxg1 neuron subsets found across multiple regions, including the medial amygdala, posterior BSTM, and parts of the preoptic region and hypothalamus; these connections could constitute a specific functional subcircuit. Supporting this, the projections of the medial amygdala reach an anterolateral hypothalamic area (Canteras et al., 1995), which includes glutamatergic neurons projecting to the basal hypothalamus (Kiss et al., 2011) and involved in inhibiting social behavior, just like those of the medial amygdala (Hong et al., 2014). All these areas contain abundant TOH-derived GFP/Foxg1 double-labeled glutamatergic cells (present results). This highlights a specific glutamatergic subcircuit from the medial amygdala to the hypothalamus involved in modulating a similar aspect of social behaviors. This subcircuit might also include GFP/Foxg1 glutamatergic neurons that spread into the pallial part of the amygdala, including the basomedial nucleus and cortical amygdalar areas that project to the medial amygdala and hypothalamus (Petrovich et al., 1996). In addition, TOH-derived neurons across these different areas could share projections to similar targets lacking GFP/Foxg1 cells, such as the septum, ventromedial hypothalamus and/or the paraventricular thalamus. Some of these projections seem plausible by studying our GFP-labeled brain material, but require experimental confirmation.

### Redefinition of SPV and contribution of its Otp-only core to the hypothalamus and adjacent regions

As noted above, only a subdivision of classical SPV, called here the SPV core, produces GFP-only cells (i.e., without Foxg1). The SPV core locates in the alar hypothalamus just ventral to the TOH. Previous studies proposed that both the complete paraventricular hypothalamic nuclear complex and the supraoptic nucleus originate in the SPV (hence the initial or classical name of this division) (Puelles and Rubenstein, 2003). Later, the proposal was refined so that the paraventricular hypothalamic complex was primarily thought to originate from the peduncular segmental division of classical SPV, while the supraoptic nuclear complex was proposed to include parts derived from the peduncular and pre-peduncular (terminal) segmental divisions of classical SPV (Puelles et al., 2012). Our data on the GFP labeling combined with Foxg1 in Otp-eGFP mouse brains throughout development contribute to further refine this proposal and help to better understand where these different nuclei originate from. Regarding the paraventricular nuclear complex, our data show that only the central and ventral parts of the paraventricular hypothalamic nucleus are rich in GFP-only cells and originate from the peduncular part of SPV core, but not the dorsal paraventricular nucleus, which is rich in GFP/Foxg1 double-labeled cells and derive from TOH.

With respect to the supraoptic nuclear complex, the vast majority of the neurons of its different subdivisions, from preoptic to tuberal levels, are GFP-only labeled (Otp-lineage), and appear to derive from SPV core. In agreement with Puelles et al. (2012), our data suggest that the supraoptic nuclear complex includes peduncular and pre-preduncular (terminal) segmental divisions. The main part of the supraoptic nucleus appears to locate at the subpial surface of the same radial domain that produces the central paraventricular nucleus, and thus appears to originate from the peduncular part of the SPV core (as noted above, this particular peduncular division can be distinguished by its high expression of Rgs4). In contrast, the part of the supraoptic nucleus that extends into the subpial surface of the ventral preoptic region appears to derive from the prepeduncular (terminal) division of SPV core. During development, cells forming this nucleus are arranged tangentially in a subpial position, and appear to extend dorsally into the TOH and beyond. Regarding the tuberal supraoptic subnucleus (named tuberal suboptic nucleus by Puelles et al., 2012, because of its position ventral to the optic chiasm), it was unclear whether its cells originate from peduncular or prepeduncular divisions of SPV (Puelles et al., 2012). In our material from E12.5 on, GFP-only cells of the tuberal supraoptic/suboptic nucleus are observed in continuation with those of the peduncular SPV core, but not with those of the terminal SPV core. This continuum of GFP-only cells is associated to the hypothalamo-hypophyseal tract, which first descents ventrally into the peduncular basal hypothalamus, and then turns rostrally into the tuberal region and the median eminence. This evidence suggests that the GFP-only cells of the tuberal supraoptic/suboptic nucleus migrate from the peduncular SPV core. In contrast to the supraoptic nuclear complex, which is rich in GFP-only labeled cells, the so-called episupraoptic nucleus (located deep to the main SO), contains a mixture of different neurons, including a subpopulation of GFP/Foxg1 double-labeled cells that appear to originate in TOH. Previous studies have suggested that the episupraoptic nucleus also contains Foxb1-expressing neurons coming from the basal hypothalamus (Alvarez-Bolado et al., 2000; Zhao et al., 2008; discussed by Puelles et al., 2012).

In addition, the SPV core also appears to produce a few GFP-only cells observed in the telencephalon (preoptic region, septum and extended amygdala), as well as a group of GFP-only cells that extends into prosomere 3 (in the prethalamus). Regarding those in the telencephalon, they represent a minor population, since the majority of the GFP labeled cells there co-express Foxg1 and appear to derive from TOH. With respect to those in the prethalamus, they seem to include a subpopulation in the zona incerta. Future studies will need to investigate the phenotype and function of these GFP-only cells.

### Foxg1-only cells in the hypothalamus

Our data show the presence of subpopulations of Foxg1-only cells (i.e. without co-expression of GFP) in the TOH mantle and the hypothalamus, including both alar and basal parts. At early embryonic stages, these cells are in continuity with those of the telencephalon, and this cellular continuum is especially evident close to the rostral end of the forebrain (i.e. along the terminal segmental division of the forebrain). Our finding agrees with data on Fogx1 expression of the Allen Developing Mouse Brain Atlas, and suggests a migration route of Foxg1 cells from dorsal to ventral in the rostral forebrain divisions. Part of these Foxg1-only cells extending into the hypothalamus may represent the migration of the gonadotropin releasing factor (GnRH)-expressing neurons, known to originate and/or migrate from the olfactory placode, and traverse the telencephalon to finally reach the hypothalamus (Forni et al., 2011a,b). Foxg1 is required for the production of mature olfactory epithelial cells from the olfactory placode, being important for maintaining the stem state of progenitors and for preventing premature differentiation of derived neurons (Duggan et al., 2008; Kawauchi et al., 2009). Foxg1 also influences GnRH cell development, since upregulation of Foxg1 expression, through depletion of specific micro RNAs (miR-9 and miR-200) involved in differentiation of olfactory receptor neurons, leads to altered genesis and positioning of GnRH neurons, resembling the problems seen in Kallmann syndrome (Garaffo et al., 2015). In addition to the possibility of the GnRH cells just mentioned, other Foxg1-only cells of the TOH and hypothalamus may originate in the telencephalon, but further investigation is required to study their exact origin and phenotype.

## Conclusions

Taken together, our results show the existence of a new radial forebrain embryonic division, the TOH, which produces GFP/Foxg1-lineage cells for a transition territory between telencephalon and hypothalamus, and includes a part of the medial extended amygdala. The TOH-derived cells of the medial extended amygdala are glutamatergic and could be engaged in a glutamatergic functional subcircuit involved in inhibiting social behavior. Since social deficits are one of the major problems seen in autism spectrum disorders (American Psychiatric Association, 2013), further studies of this neuron subpopulation in different animals, including humans, become essential. Future studies should also evaluate the implication of the different glutamatergic and GABAergic neurons of the extended amygdala in the excitatory/inhibitory neuron imbalance reported in several neurodevelopmental and mental disorders (Mariani et al., 2015; Selten et al., 2018; Fogaça and Duman, 2019). This imbalance is often analyzed by only focusing on the cortex, but such studies should also include other areas known to be affected in these disorders such as the amygdala.

Different developmental-based observations reveal the high complexity of brain cellular architecture, and raise questions on the mature molecular phenotype of each neuron subpopulation, on their integration in different functional networks, and on the interaction between different subtypes. These studies can clearly help to a better understanding of functional subcircuits based on developmental results, aiming to fill the gap between genes, cells, circuits and functions (Sokolowski and Corbin, 2012; Medina et al., 2011; 2017a,b), and might provide baseline data for future, more precise circuit-based approaches to psychiatry (Lisman et al., 2008; Spellman and Gordon, 2015; Gordon, 2016).

## Acknowledgements

Supported by grants from the Spanish Ministerio de Economía y Competitividad (MINECO) and Fondo Europeo de Desarrollo Regional (FEDER) (ref. BFU2015-68537-R) and Ministerio de Ciencia e Innovación (ref. PID2019-108725RB-100). Lorena Morales had a predoctoral fellowship from Universitat de Lleida i Ajuts Jade Plus, and a predoctoral contract from IRBLleida/Diputació de Lleida.

## Availability of data

Data available on request from the authors. The data that support the findings of this study are available from the corresponding author upon reasonable request.

## Abbreviations

AAv: ventral anterior amygdala
ac: anterior commissure
AH: anterior hypothalamus
BC: amygdalar basolateral complex
BMa: basomedial amygdala, anterior subnucleus
BSTM: medial bed nucleus of the stria terminalis
BSTMa: BSTM, anterior division
BSTMh: BSTM, hypothalamic subdivision/ hypothalamic subdivision of BSTM
BSTMp: BSTM, posterior division
BSTL: lateral bed nucleus of the stria terminalis
Ce: central nucleus of the amygdala (or simply central amygdala)
Cer: cerebellum
CPu: caudate-putamen complex
DM: dorsomedial hypothalamic nucleus
EM: median eminence
ESO: episupraoptic nucleus
f: fornix
GP: globus pallidus
HF: hippocampal formation
Hya: alar hypothalamus
Hyb: basal hypothalamus
LH: lateral hypothalamus
LH: basal LH
LOT: nucleus of the lateral olfactory tract
LPO: lateral preoptic area
lv: lateral ventricle
m: mantle
Ma: mammillary nuclei
Me: medial nucleus of the amygdala (or simply, medial amygdala)
MeA: anterior subnucleus of Me
MeP: posterior subnucleus of Me
MePD: dorsal subnucleus of MeP
MePV: ventral subnucleus of MeP
MePVm: medial branch of MePV
MePVl: lateral branch of MePV
MGEvc: medial glaglionic eminence, ventrocaudal subdivision
NH: neurohypophysis
OB: olfactory bulb
oc: optic chiasm
op: optic pedicle
ot: optic tract
P: pallium
Pad: paraventricular nucleus, dorsal subdivision
Pav: paraventricular nucleus, ventral subdivision
Pt: pretectum
PHy: peduncular hypothalamus
POPe: periventricular part of the preoptic area
POs: subpallial preoptic area
POv: ventral preoptic area
PTh: prethalamus
PThE: prethalamic eminence
PVN: paraventricular nucleus, principal subdivision
Se: septum
SEA: sublenticular extended amygdala
SO: supraoptic nucleus, principal subdivision
SOt: supraoptic nucleus, terminal subdivision
SOtu: of the supraoptic nucleus, tuberal subdivision
Sp: subpallium
SC: superior colliculus
SCN: suprachiasmatic nucleus
SPa: subparaventricular region
SPV: supraopto-paraventricular hypothalamic domain (classical definition)
SPVc: supraopto-paraventricular hypothalamic domain, core part
svz: subventricular zone
Tel: telencephalon
Th: thalamus
THy: terminal hypothalamus
TOH: telencephalo-opto-hypothalamic embryonic domain
VMH: ventromedial hypothalamic nucleus
vz: ventricular zone
3v: third ventricle
4v: fourth ventricle

